# Age-Related Differences in Affective Behaviors in Mice: Possible Role of Prefrontal Cortical-Hippocampal Functional Connectivity and Metabolomic Profiles

**DOI:** 10.1101/2023.11.13.566691

**Authors:** Marcelo Febo, Rohit Mahar, Nicholas A. Rodriguez, Joy Buraima, Marjory Pompilus, Aeja M. Pinto, Matteo M. Grudny, Adriaan W. Bruijnzeel, Matthew E. Merritt

## Abstract

The differential expression of emotional reactivity from early to late adulthood may involve maturation of prefrontal cortical responses to negative valence stimuli. In mice, age-related changes in affective behaviors have been reported, but the functional neural circuitry warrants further investigation. We assessed age variations in affective behaviors and functional connectivity in male and female C57BL6/J mice. Mice aged 10, 30 and 60 weeks (wo) were tested over 8 weeks for open field activity, sucrose preference, social interactions, fear conditioning, and functional neuroimaging. Prefrontal cortical and hippocampal tissues were excised for metabolomics. Our results indicate that young and old mice differ significantly in affective behavioral, functional connectome and prefrontal cortical-hippocampal metabolome. Young mice show a greater responsivity to novel environmental and social stimuli compared to older mice. Conversely, late middle-aged mice (60wo group) display variable patterns of fear conditioning and with re-testing with a modified context. Functional connectivity between a temporal cortical/auditory cortex network and subregions of the anterior cingulate cortex and ventral hippocampus, and a greater network modularity and assortative mixing of nodes was stronger in young versus older adult mice. Metabolome analyses identified differences in several essential amino acids between 10wo mice and the other age groups. The results support differential expression of ‘emotionality’ across distinct stages of the mouse lifespan involving greater prefrontal-hippocampal connectivity and neurochemistry.

## Introduction

The U.S. is seeing a rise in affective conditions among young adults (Kessler et al., 2012;Terlizzi and Villarroel, 2020). The most recent national mental health statistics show that young adults (ages 18-25 years) have a greater prevalence of anxiety and mood conditions but seek less treatment than older and middle-aged adults (ages 50 and older) (SAMHSA, 2022). Prior studies have revealed age-related changes in ‘emotionality’ and emotion processing that may contribute to a differential vulnerability of young individuals to experience an affective condition. Compared to older individuals, young adults cite greater negative emotional experiences, show less emotional control (Gross et al., 1997) and stability (Carstensen et al., 2011), have a greater tendency to report negative mood states (Machado et al., 2019), more worries (Brenes, 2006), and a greater recall of negative memories (Charles et al., 2003). Neuroimaging studies have determined the major components of neural circuitry involved in the differential processing of negative valence stimuli between young and older adults. The dorsal and rostral subregions of anterior cingulate cortex (ACg) show weaker responses to negative stimuli in old versus young adults (Chaudhary et al., 2023). Functional magnetic resonance imaging (fMRI) studies employing face perception tasks indicate that the amygdala has greater blood oxygenation level dependent (BOLD) response to negative emotion stimuli in young compared older adults (Iidaka et al., 2002;Fischer et al., 2005;Tessitore et al., 2005). Other studies, however, observed either no age associated differences in the amygdala response to negative stimuli (Wright et al., 2006) or report an increased amygdala functional connectivity from adolescence to adulthood (Ravindranath et al., 2020). Recent evidence suggests that the neurobiological adaptations associated with adulthood variations in affective behaviors likely involves a larger number of brain areas outside the amygdala and changes in neural interactions at a network level (Malezieux et al., 2023;Wang et al., 2023).

Age-related changes in affective behaviors in early versus late adulthood have been reported in mice (Flurkey et al., 2007;Shoji and Miyakawa, 2019;Baumgartner et al., 2023). Young (2–3-month-old; mo) C57BL/6J male mice display greater anxiety-like behaviors compared to older adult, late middle age and aged mice (Ammassari-Teule et al., 1994;Shoji and Miyakawa, 2019). Although this remains unclear, since other studies provide evidence of greater anxiety-like behaviors in aged compared to young adult mice (Shoji and Miyakawa, 2019;Baumgartner et al., 2023). Nevertheless, the expression of fear-related associative memory (von Bohlen und Halbach et al., 2006;Evans et al., 2021;Ferreira et al., 2023), reward sensitivity (Ammassari-Teule et al., 1994;Malatynska et al., 2012), sociability and social recognition (Shoji and Miyakawa, 2019), and learned helplessness on a Porsolt test (Ferreira et al., 2023) are all observed to vary in young versus older adulthood mice. In parallel with these affective behavior findings, there is recent evidence of brain structural differences between early and late adulthood (Clifford et al., 2023). In an analysis of structural covariance network data, it was shown that young mice have a higher modularity with greater node degree in anterior cingulate cortex and lower in hippocampal formation than older adult mice (Clifford et al., 2023). This age-related difference in structural network modularity was linked to cognitive differences (Clifford et al., 2023). It remains unclear whether functional network topology differs in a similar manner to the reported structural changes in young versus older adult mice. The relationship between functional network activity, as measured by functional magnetic resonance imaging (fMRI), and affective behaviors in early and late adulthood warrants investigation. In the present study, we measured affective behaviors involving anxiety-like open field activity, reward sensitivity, sociability, and fear memory in young adult, older adult, and late middle aged C57BL/6J male and female mice. We also evaluated the relationship between behavioral measures and fMRI markers of network connectivity. Finally, we conducted hippocampal and prefrontal cortical metabolomic analysis and observed several novel protein-related substrates differing between young, older adult and late middle-aged mice.

## Materials and Methods

### Subjects

Male and female C57BL/6J (B6) mice aged 10, 30, or 60 weeks old (wo) at the time of arrival were allowed a week of acclimation to the vivarium prior to the start of experiments (N=30, total; n=10 per age group, 50% females; The Jackson Laboratory, Bar Harbor, ME). Female mice initially weighed (in grams; mean ± standard deviation) 19±0.93, 32±4.6, 29±2.8 and males 26.7±2, 30±2.5, 46.8±5.6 (for 10, 30, 60wo, respectively). The selected age groups represent 3 stages of adulthood. B6 mice at 3-6MO (12-24wo) are considered mature adults and at 10-14MO (40-56wo) are considered middle age (Flurkey et al., 2007). The youngest 10wo group (2.5MO) was tested just before mature adulthood and the 60wo group was just at the end of middle age (15MO). At 30wo (7.5MO), mice start to show early signs of senescence (such as retirement from breeding at 8MO) and were included here as reference to compare early adulthood and late middle age changes (Flurkey et al., 2007). Mice were housed in age- and sex-matched groups of 3-4 in a temperature- and humidity-controlled room, inside conventional air filtered cages (dimensions: 29 x 18 x 13 cm) with food and water available *ad libitum* (vivarium lights on from 07:00-19:00 hours). All procedures received prior approval from the Institutional Animal Care and Use Committee of the University of Florida and follow all applicable NIH guidelines.

### Experimental design

**Figure 1** shows the experimental timeline. All behavioral tests and functional neuroimaging were conducted over an 8-week period. Behavioral tests included locomotor activity in a novel environment, sociability and social recognition tests, sucrose preference tests, and cued fear conditioning (**Figure 1**). Functional magnetic resonance imaging (fMRI) was conducted a week after sucrose preference testing and two weeks before the final behavioral test (fear conditioning). At the end of the experiments, mice were deeply anesthetized with 5% isoflurane, and brain tissue harvested for the metabolomic assay of excised prefrontal cortex and hippocampus tissue. Imaging was thus carried out when mice were 14, 34 and 63wo and these were euthanized 8 weeks after arrival, at 18, 38 and 68wo. However, to simplify presentation of the results, we keep group names in figures as 10wo, 30wo and 60wo.

**Figure 1.**
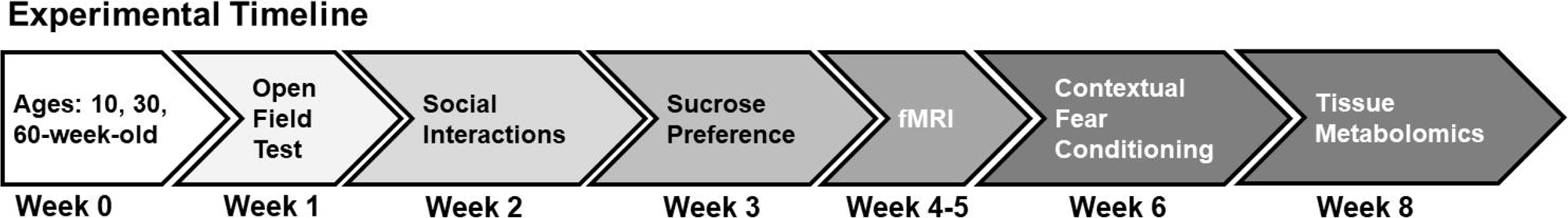
Experimental timeline. See text for details.

### Locomotor activity in a novel environment

An automated rodent activity monitoring cage system (Accuscan Instruments, Columbus, OH) was used to measure spontaneous locomotor activity in a novel environment (Bruijnzeel et al., 2016). The day before locomotor measurements, mice were acclimatized for 1 hour to the testing room. The activity monitors are made of clear Plexiglas (40 cm^3^) with an outer steel frame containing infrared emitting and receiving panels. Each panel has 16 equally spaced (2.5 cm) infrared beams crossing the length and width of the cages at a height of 2 cm (horizontal beams). Several patterns of sequential and repetitive infrared beam interruptions were registered and sent to a programmable counter, which sorted beam counts and sent sorted data to a PC running Versamax™ software. The sorted activity counts were classified as sequential beam breaks (forward locomotion), repetitive beam breaks (putative repetitive behaviors), total, center and margin distances travelled (in cm). Activity was measured during a single 30-minute test.

### Sucrose preference test

The sucrose preference test was conducted as described previously, with modifications (Bruijnzeel et al., 2016;Chellian et al., 2020). A decrease in preference to consume a 2% sucrose solution over water suggests anhedonia (D’Aquila et al., 1997). ‘Leakproof’ bottles (100 ml, graduated by 1 ml) with stoppers and sipper tubes were purchased (Braintree Scientific Inc, Braintree, MA; model no. WTRBTL). Filtered home cage lids with two bottle openings (Allentown, Inc., Allentown, NJ), were used for two bottle fluid delivery in mouse home cages. Mice were temporarily single housed for testing and once testing concluded they were returned to their original cages. First, mice were acclimatized to bottles filled with 2% sucrose solution for 3 days (96 h). On each of the 3 days, bottles were weighed (in grams) to monitor fluid intake and replaced with a full bottle. The 96h acclimation period was followed by a single 24 h two-bottle choice test in which one bottle was filled with 2% sucrose solution and the other with water. The side of the cage lid where the sucrose and water filled bottles were placed was counterbalanced across animals. Water and sucrose consumed was measured after a 24 h period (McCall et al., 2013). Bottles were weighted and values converted from grams to ml. An index (S_I_) of how much sucrose (s) consumed versus water (w) was calculated as: 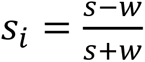.

### Sociability and social recognition test

We used a modified version of a previously reported sociability and social recognition test protocol (Tan et al., 2019). Test arenas were made of clear Plexiglas, divided by walls into 3 equal sized quadrants (arena total length x width x height dimensions: 60 x 45 x 20 cm). The middle quadrant is the neutral zone, and the side quadrants (left and right) were used for the placement of ‘bait’ mice for sociability and recognition tests. Each side quadrant can be entered through a central door in the dividing walls. A pair of clear plastic ‘bait’ cups covered with equally sized and equidistant perforations for air flow were inverted over the ‘bait’ mouse for sociability testing. Test mice were acclimatized to the room and the neutral quadrant of the test arena on 3 days, 15 minutes a day. During sociability testing, a novel (unfamiliar) mouse (an adult mouse of the same strain and sex; age range 3-6mo) was placed under a cup and the entrance to the ‘bait’ side of the arena was opened for exploration by the test mouse while the other side is kept sealed. In preliminary tests, this encouraged mice to explore the unfamiliar ‘bait’ mouse more vigorously during a first encounter. Mice were tested for 10 minutes for time spent exploring the unfamiliar mouse. Upon completion of the first test, and after a 5-minute break to clean the arena, mice were placed back into the arena in the presence of a novel mouse and the previously encountered (now familiar) mouse and re-tested for 10 more minutes with both sides opened for exploration. In the first test, the degree of exploration of an unfamiliar mouse (“novel 1”) is considered here as a measure of the degree of sociability. In the second test, less exploration of the familiar mouse (“familiar 1”) relative to the new unfamiliar mouse (“novel 2”) is a sign of social recognition (Crawley et al., 2007). Behaviors were recorded using a digital camera directly connected via FireWire to a Linux computer. We used the ‘*dvgrab*’ tool from the Unix command terminal to program video capture settings and store the recorded videos.

Behavioral analyses were carried out using DeepLabCut (DLC, version 2.2.1) (Mathis et al., 2018;Nath et al., 2019). We labeled 120 frames of mouse trajectories within the test cages taken from 6 different videos and 95% of those labels were used to train a machine learning algorithm. A ResNet-50-based neural network with default parameters for 2 training iterations was used for training. Data (digital videos) were split into training and test datasets. The training dataset was used to train a DLC model. Conversely, the test dataset was used to evaluate the performance of the model. We validated the DLC model with 7 train-test-split shuffles. The model with the smallest mean average Euclidean error (MAE) was selected. Before model selection, we applied a p-value cutoff of 0.6 to exclude model predictions with a likelihood under 0.6. The final selected model had a MAE of 2.9 pixels. This neural network was used to analyze videos from similar experimental settings to generate trajectory data. Using the R packages ‘trajr’ (Mclean and Volponi, 2018) and ‘DLCanalyzer’ (Sturman et al., 2020), spurious trajectories were removed before analyses. The videos were analyzed in two parts, the first sociability test (T1) and the second social recognition test (T2). The immediate area surrounding and including the mouse ‘bait’ cup on one side of the arena was designated ‘ROI A’ and the other ‘ROI B’ (region of interest A and B, respectively). A python script was compiled to generate the number of entrances, the velocity (in pixels per frame or ppf), and the duration of initial approach and time spent within the ROIs.

### Classical fear conditioning

Fear conditioning was conducted two weeks following the fMRI studies. The fear conditioning cages were housed in sound attenuation chambers. The chambers had internal computer-controlled lighting, white noise generator, a fan, and control units for distinct aspects of the fear conditioning assay. These included audio speaker controls for tone generation, shock controllers for the internal cage floors, and infrared and white lighting controllers. The internal operant cage was made of translucent Plexiglas with a steel frame and steel shock grid floor connected to a computer controlled current stimulator (Med-Associates, Inc. St. Albans, VT). Activity within the fear conditioning cages was captured by a monochrome video camera and video recordings were optimized via externally controlled near infrared (NIR), white light and sound controllers. All cage accessories were controlled by PC running Video Freeze Software^TM^. Camera lens brightness, gain, and shutter settings were verified and adjusted before collecting data and were kept constant in all mice. For each session, the detected motion-sensitive signals were calibrated with the mouse inside the cage and NIR lights on. The estimated motion index was set to the same threshold level (10 a.u.) in all mice prior to data exporting or analyses.

Videos were acquired at 30 frames per second (fps) and the minimum freeze duration was 30 frames. For training on the first day, a 180 second baseline was followed by four presentations of a 20 second tone (90 dB, 5kHz, 50ms risetime) with the presentation of moderate level current to the entire grid floor for 1 second (0.9mA) (inescapable shock) on the final second before the tone ended. This 90dB tone is above the auditory brainstem response measured in C57BL/6J mice, which declines with age and reaches 60-80dB at 5-8kHz (Henry, 2004;Yu et al., 2011;Youn et al., 2020). Each of the four tone-shock presentations were separated by 190 seconds and the entire session lasted 17 minutes. On days 2 and 3 (recall tests at 24 and 48 h of training day 1, respectively), the same protocol was carried out but in the absence of a shock. On day 3 (modified context test), the cage environment was modified by placing a plastic smooth white floor and diagonal walls, which covered the grid floor, light, and speaker.

For each of the time epochs corresponding to the 4 tone-shock pairings, a 20 second pre-shock tone interval and a 40 second post-shock interval were used to quantify number of freezing bouts (freezing counts) in a total 60 second epoch (per day there were four 60 second epochs summed to provide total freezing counts per day). Motion-index was also used as a secondary measure of general locomotor activity inside the cage during the same 60 second epochs/day.

### Magnetic resonance imaging instrumentation

Images were collected on a magnetic resonance spectrometer tuned to 200.6 MHz proton resonant frequency (4.7 Tesla/33 cm actively shielded magnet). The MR system was equipped with Resonance Research Inc. spatial encoding gradients (RRI BFG-200/115-S14, maximum gradient strength of 670 mT/m at 300 Amps and a 120 µs risetime; RRI, Billerica, MA) and controlled by a VnmrJ 3.1 console (Agilent, Palo Alto, CA). A custom-built 8.8 cm x 11.6 cm radiofrequency (RF) transmit coil used in combination with a 3 cm x 3.5 cm actively decoupled/tunable RF receive quadrature surface coil (RF engineering lab, Advanced Magnetic Resonance Imaging and Spectroscopy Facility, Gainesville, FL).

### Mouse imaging setup

Mice were scanned sedated under a continuous paranasal flow of 0.25 % isoflurane gas (delivered at 0.04L/min mixed with medical grade air containing 70% nitrogen and 30% oxygen, Airgas, Inc.) and a continuous subcutaneous infusion of dexmedetomidine. Prior to setup, mice were anesthetized under 2% isoflurane and administered an intraperitoneal (i.p.) injection of 0.1mg/kg dexmedetomidine (at a 1ml/kg volume). Isoflurane was reduced to 0.25% throughout the remaining imaging session. An infusion line was setup for subcutaneous delivery of dexmedetomidine over the course of scanning (0.1mg/kg/ml at an infusion rate of 25 µl/h using a PHD-Ultra microinfusion pump, Harvard Apparatus, Holliston, MA). Functional MRI scans were collected at least 50 minutes after the i.p. injection. Respiratory rates were monitored continuously, and body temperature was maintained at 36-37°C using a warm water recirculation system (SA Instruments, Inc., New York).

### Functional MRI acquisition protocol

We acquired one T2 weighted anatomical and an fMRI scan per mouse. The T2-weighted fast spin echo (FSE) sequence was acquired with the following parameters: effective echo time = 48 ms, repetition time (TR) = 2 seconds, echo train length = 8, number of averages = 12, field of view (FOV) of 19.2 mm x 19.2 mm and 0.8 mm thick slice, and a data matrix of 192 x 192 (0.1 µm^2^ in plane) and 12 interleaved ascending coronal (axial) slices covering the entire brain from the rostral-most extent of the anterior prefrontal cortical surface, caudally towards the upper brainstem and cerebellum (acquisition time 9 minutes 36 seconds). Functional images were collected using a single-shot gradient echo planar imaging sequence with the following parameters: TE = 22 ms, flip angle = 90°, TR = 1.2 seconds, 600 repetitions, FOV = 19.2 x 19.2 mm and 0.8 mm thick slice, and a data matrix of 48 x 48 (0.4 µm^2^ in plane) with 12 interleaved ascending coronal slices in the same position as the anatomical scan. Ten dummy EPI scans were run prior to acquiring data under steady state conditions. Respiratory rates, isoflurane and dexmedetomidine delivery, body temperature, lighting, and room conditions were kept constant across subjects.

### Image pre-processing

The image processing workflow is illustrated in **Figure 2** (Colon-Perez et al., 2019;Kotlarz et al., 2022;Sadaka et al., 2023;Sakthivel et al., 2023). Resting state processing was carried out using Analysis of Functional NeuroImages (AFNI) (Cox, 1996), FMRIB Software Library (FSL) (Jenkinson et al., 2002), and Advanced Normalization Tools (ANTs) (Klein et al., 2009). Binary masks for anatomical and functional scans were created using ITKSNAP (Yushkevich et al., 2006). The brain binary masks were used for brain extraction prior to registration steps. Times series spikes were removed (3dDespike, AFNI), image repetition frames aligned to the first time series volume (3dvolreg, AFNI), and detrended (high pass filter <0.009 Hz using AFNI 3dTproject). Independent component analysis (ICA) using Multivariate Exploratory Optimized Decomposition into Independent Components (FSL MELODIC version 3.0) was used to assess structured ‘noise’ or artefact components in each subject scan, in their native space. Most, if not all ICA components in this first stage contained artefact signal voxels along brain edges, in ventricles, and large vessel regions. These components were suppressed using a soft (‘non-aggressive’) regression approach, as implemented in FMRIB Software Library (FSL 6.0.3) using fsl_regfilt (Jenkinson et al., 2002). A low-pass filter (>0.12Hz) and spatial smoothing (0.4mm FWHM) was next applied to the fMRI scans prior to registration steps. Post-regression ICA was carried out to verify removal of artefact components and preliminary assessment of putative ICA networks in individual scans.

**Figure 2.**
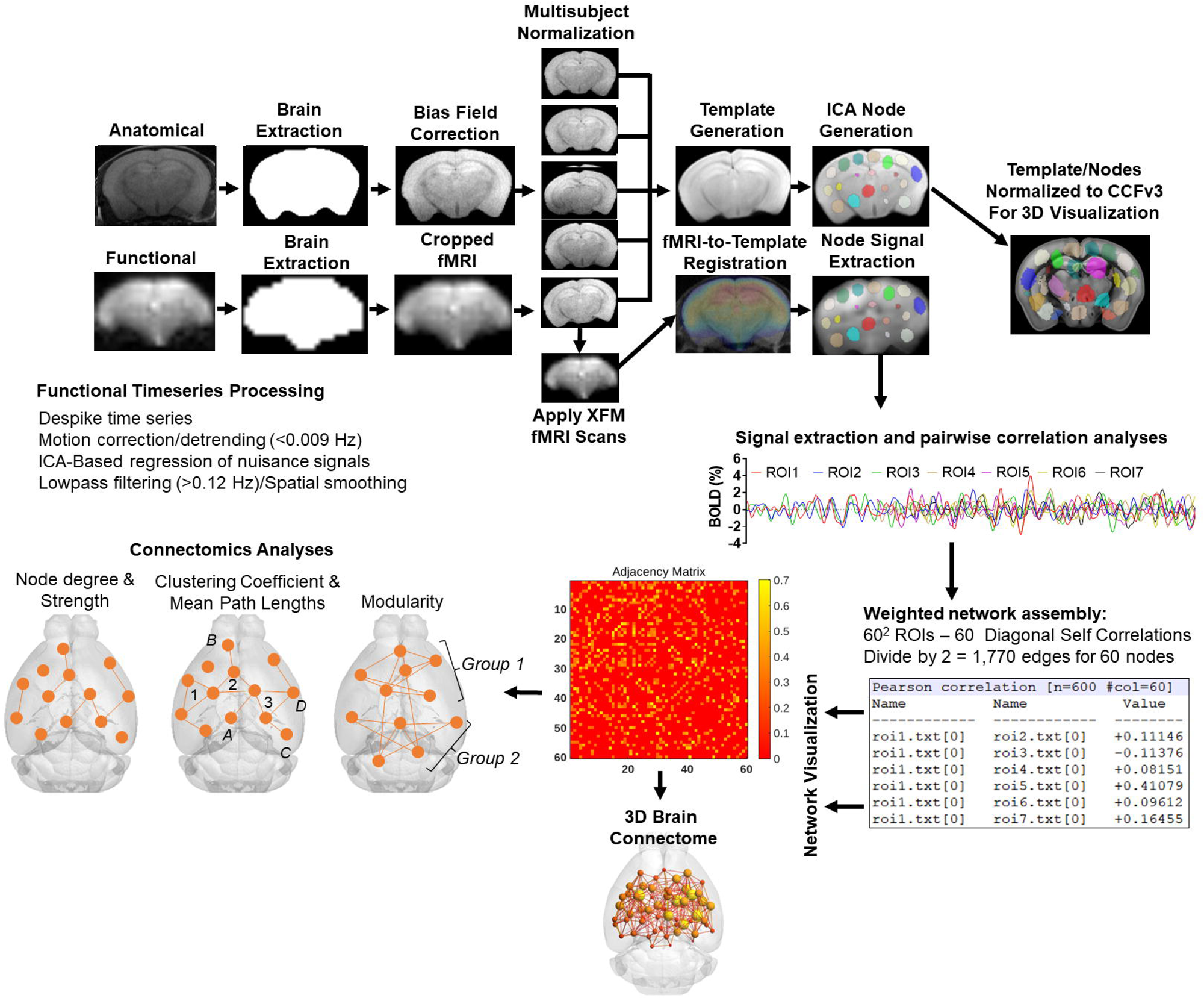
Image processing workflow. Anatomical scans from all subjects were used to generate a group-wise template for anatomical and functional image registrations and individual and group ICA-based signal processing steps. The decomposition of whole brain signals into 60 components was used to generate region of interest masks for signal extraction and pairwise correlations. The study specific node-set were normalized to the common coordinate framework version 3 mouse atlas for 3D visualizations of nodes and edges. See text for details.

### Image template construction, subject normalization, and resting state signal extraction

Anatomical scans were cropped, and bias field corrected (N4BiasFieldCorrection, ANTs). Functional scans were cropped, and a temporal mean image was registered to the anatomical. The functional-to-anatomical registration matrix was applied to all fMRI volumes. A multi-T2 anatomical template was generated using ANTs ‘antsMultivariateTemplateConstruction2.sh’ script (cross correlation used as a similarity metric and a greedy SyN as transformation model). Individual subject-to-template transformation matrices were applied to fMRI datasets. A group ICA was applied to template-registered concatenated fMRI scans to generate a set of 60 nodes for ROI-based intrinsic fMRI signal extraction, pairwise cross-correlation statistics, and adjacency matrix construction. The ICA was set to an upper limit 60 component signal decomposition and a mixture model threshold of 0.7. The resulting 60 ICA components included peak signal (peak T statistic) nodes in midline prefrontal cortex (anterior cingulate), somatosensory regions, temporal, visual, auditory cortices, entorhinal cortex, several hippocampal nodes, subiculum, amygdala, thalamic nuclei, and striatum. Each of the 60 nodes were binarized and assessed to ensure non-overlapping coverage of brain areas on the template and registered scans (**Figure 2**). Both the template and ICA nodes were normalized to the Allen Brain Institutes common coordinate framework version 3 (CCFv3) atlas of the B6 mouse brain for final 3D visualizations (Lein et al., 2007). Signal timeseries were extracted from preprocessed fMRI scans with the assistance of 60 ROI mask overlays. This generated 60 individual ROI text files per subject that contained L2-normalized resting state signals as a vector of 600 data points.

The timeseries files were used in cross-correlations and in calculations of Pearson r coefficients for every pairwise combinations of ROIs (using *corrcoef* function in MATLAB). The resulting number of pairwise correlations was 1,710 per subject (after removing 60 self-correlations). Correlation coefficients were Fisher’s transform to ensure normality prior to statistical analyses. The center voxel coordinates for the ICA-based nodes normalized to the CCFv3 were used in 3D network visualizations using BrainNet viewer (Xia et al., 2013).

### Network analysis

Weighted undirected matrices were analyzed using Brain Connectivity Toolbox (Rubinov and Sporns, 2010), igraph (version 1.5.1.9005) (G. and Nepusz, 2006) in R, and tools in MATLAB (Mathworks, Natick, MA). Global graph metrics were calculated for edge density thresholds ranging from 6-32%. Global network measures for this density range were converted to area under the curve (AUC) values prior to multivariate statistical assessments. Node-specific (ROI) network measures were calculated at a 16% threshold, which in the present dataset corresponds to a Pearson r of approximately 0.2 or more, an average degree of 9.1, and network strength of 2.5-5.0 (**Supplemental Figure 1**). As a comparison, graph density based on bootstrapped calculations of confidence intervals for significant Pearson correlations corresponds to a significant r of ≥ 0.06, 40% density (δ ≥ 0.4), degree(k) ≥ 20, and strength ≥ 4 (**Supplemental Figure 1**).

A major objective of this study was to assess maturational and aging changes in functional network topology and determine links between network measures and spontaneous ‘affective’ behaviors. To this end, we assessed several graph measures of network integration and communication efficiency. These included node clustering coefficient (CC; number of interconnected node pairs connected to a node) (Onnela et al., 2005), node strength (sum of edge weights/node) and degree (sum of edges/node) and global measures, such as transitivity (related to CC; number of triad groups normalized by all possible triad nodes in a network), mean distance or characteristic path length (CPL; the average edges or edge weights between node pairs), the small world coefficient (SWC) (Humphries et al., 2006;Humphries and Gurney, 2008) and small world measure (SWM) (Telesford et al., 2011). Global efficiency is the inverse of CPL. For local node efficiency, a length matrix was used to calculate inverse values in vicinity of a node, with added weights used to emphasize the highest efficiency node paths (Rubinov and Sporns, 2010;Wang et al., 2017). To corroborate results relative to random networks, all network measures were calculated on original and randomized versions of the functional connectivity matrices. Positive and negative edges were randomized by ∼5 binary swaps with edge weights re-sorted at each step. The approach preserves weights, degree, and strength distributions (Rubinov and Sporns, 2010). We confirmed that the randomization approach preserved weights per nodes, node degree and strength distributions (**Supplemental Figure 2**). Node strength and degree distributions were similar between random and original networks, with distributions having long tail at low density threshold and near Gaussian at high density (**Supplemental Figure 2A-F**). Degree and strength were approximately preserved across nodes, although the specific organization of edge weights differed when viewed on 3D brains (**Supplemental Figure 2G-K**). The positive and negative matrix strength sequence accuracies for input and output matrices were r = 0.99 and r = 0.98. SWC was calculated as the ratio of gamma to lambda coefficients, where gamma is the ratio of real to random network CC, and lambda the ratio of real to random network CPL (Humphries and Gurney, 2008). SWM was calculated with an artificial non-ring lattice network as reference, as previously reported (Telesford et al., 2011).

A probabilistic approach for community detection was used to calculate a modularity statistic (Q), which indexes the rate of intra-group connections versus connections due to chance (Blondel et al., 2008). The procedure starts with a random grouping of nodes and iteratively moving nodes into groups which maximize the value of Q. The final number of modules and node assignments to each group (e.g., community affiliation assignments) was taken as the median of 100 iterations of the modularity maximization procedure (Pompilus et al., 2020). The number of communities per mouse brain functional network and population size of each community were also assessed. We evaluated other community detection routines in igraph as a comparison to results obtained with the implementation of Louvain’s algorithm in Brain Connectivity Toolbox (results in **Table 1**). The role each node plays within their assigned module can vary according to their participation coefficient. The participation coefficient is calculated as: 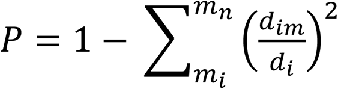, where *d_im_* is the within-module node degree and *d_i_* is the overall network degree for a node.

**Table 1.**
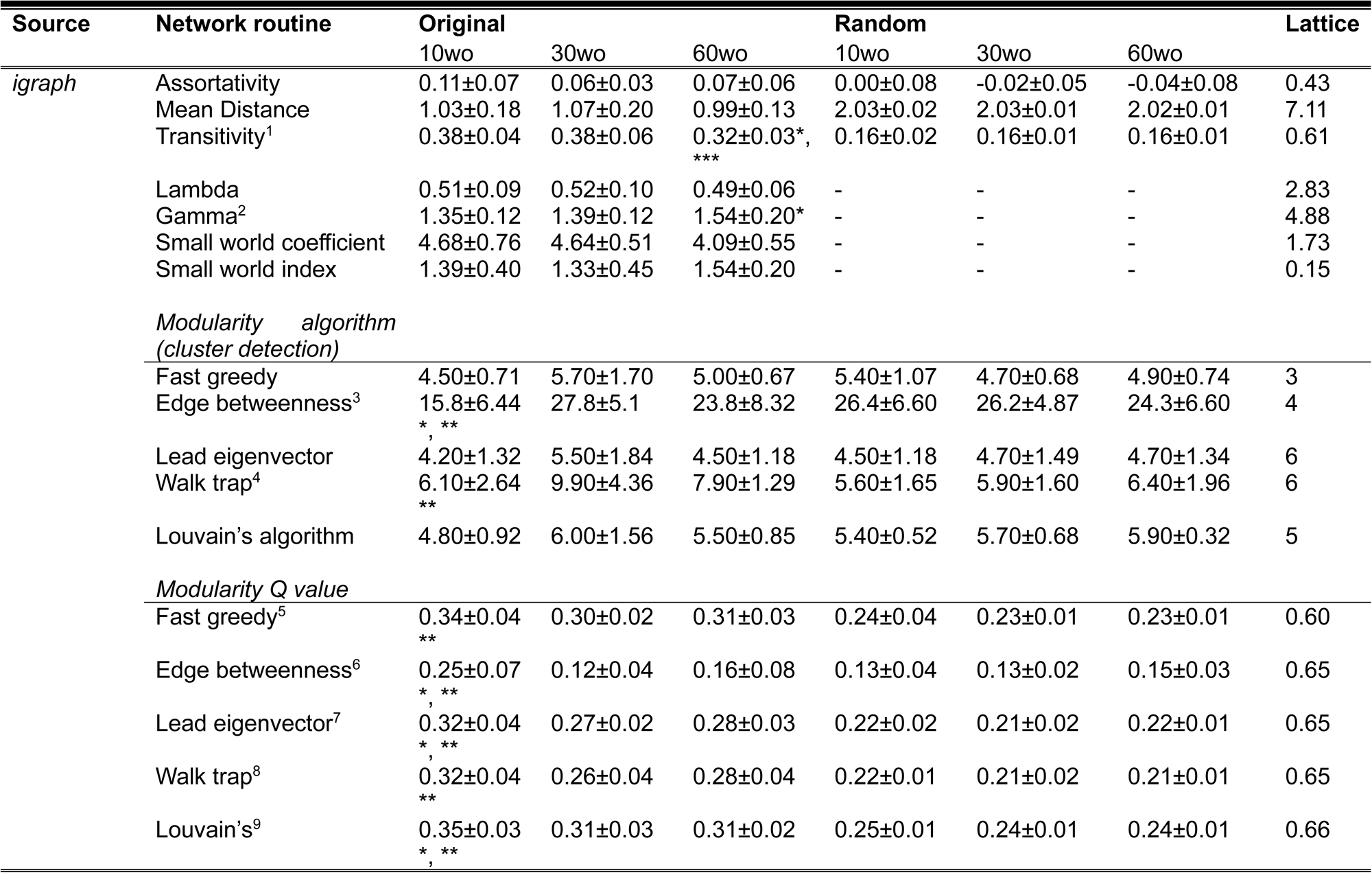
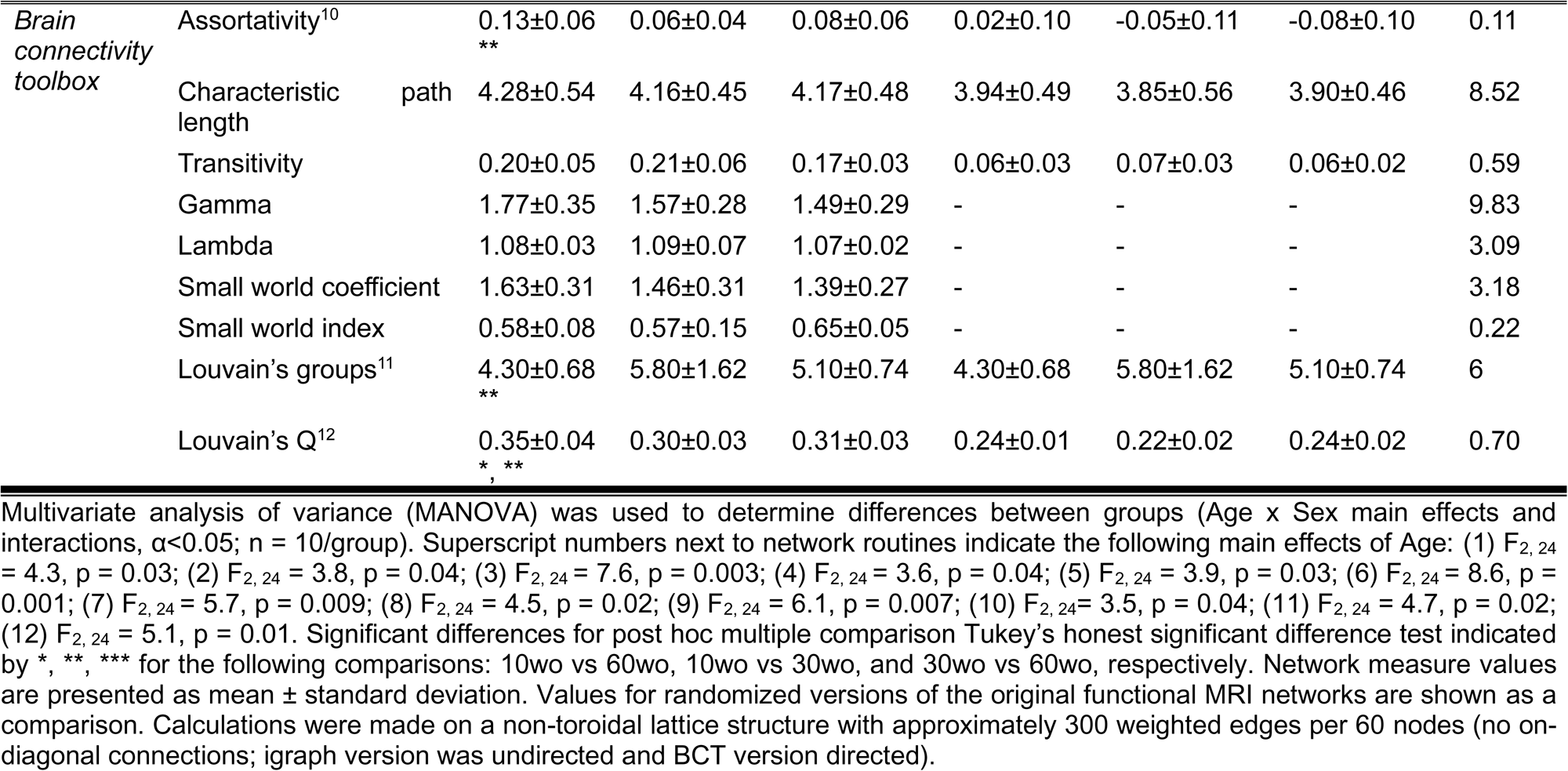
Age differences in modularity persist across different cluster detection methods. All network measures calculated on mouse functional connectivity graphs with an edge density threshold preserving the top 16% Pearson correlations. Randomization of original networks preserved weights, node degree and strength sequences. Synthetic lattice structure used in small world index calculation was generated with the same number of nodes and approximately similar edge density.

We analyzed the tendency of assortative vs dissortative mixing of nodes (Newman, 2002). The assortativity index is a correlation coefficient comparing node strength values between pairs of edge-connected nodes. Positive r values indicate connectivity between pairs of nodes with similar strengths (e.g., high strength nodes pairs with high and low with low), while negative r values indicate cross-pairings between low and high strength nodes. We generated and analyzed weighted rich club coefficient curves (*Φ*) at an edge density of 16% (Colizza et al., 2006). Nodes belonging to a rich club subnetwork have an above-chance tendency to tightly connect with each other and share the bulk of connections within the network (Colizza et al., 2006;van den Heuvel and Sporns, 2011). The approach creates subgraphs containing nodes with strength values at a predetermined degree value, *k*. For each k-level subgraph, *Φ_w_* is calculated as the ratio of the sum of edge weights that connect nodes of degree k or higher (*W_>k_*) to the total possible number of summed edge-weights (van den Heuvel and Sporns, 2011). Rich club coefficient values were calculated for original and randomized networks and a normalized rich club index analyzed (i.e., (*Φnorm = Φorig/Φrand*).

Functional connectivity networks were visualized in BrainNet viewer (Xia et al., 2013). Center coordinates for each node were based on CCFv3 parcellations, as indicated above. Several 3D mouse brain connectome maps were generated in which size of nodes (spheres) was scaled according to AUC values for several node-centric measures at various thresholds. Additional 3D maps were generated with node colors representing the module assignment (e.g., community affiliation vector) and node size weighted by modularity index.

### Sample Preparation for Gas Chromatography-Mass Spectrometry (GC-MS)

Fresh brain tissue was extracted, and two isolated regions were weighed, and flash frozen for metabolomics (**Supplemental Figure 3**). A 1mm thick slice of the prefrontal cortex was manually dissected from approximately Bregma anterior-posterior −2.96mm to −1.94mm (Paxinos and Franklin, 2001), and a 2mm wide midline cut including from the median −/+ 1mm medial lateral structures −3 to −3.5 mm deep. The dissected regions thus included areas of the infralimbic, prelimbic, medial orbital, anterior cingulate and midline primary motor cortices. The dorsal hippocampus was dissected at −1.46 mm rostrally to about −2.46 mm caudally (Bregma coordinates) and stripped of its overlying cortex and white matter. These regions have been implicated previously in age related differences in structural connectivity (Clifford et al., 2023).

Mice brain tissue (20-25 mg) was extracted with 1 mL of Acetonitrile:Isopropanol:Water (3:3:2, *v*:*v*:*v*), centrifuged for 25 min at 10,000× *g*. The supernatant was transferred into a new tube and dried in a speedvac. The dried extract was resuspended in 0.5 mL of Acetonitrile:Water (1:1, *v*:*v*) and centrifuged for 15 min and the supernatant was transferred into reaction vial and dried down under nitrogen. Fifty microliters of 2% MOX reagent were added to the vial and incubated at 30 °C for 90 min followed by addition of 50µL of MTBSTFA +TBDMS and incubation at 60 °C for 1 h. Finally, reaction solution was transferred into GC vial for GC-MS analysis (Mahar et al., 2021).

### GC-MS analysis

GC-MS data was recorded with Single Quadrupole Mass Spectrometer and GC (Trace 1310) (Thermo Fisher Scientific, USA). The column was 30 m long dimethyl (95%)/diphenyl polysiloxane (5%) RTX-5MS with 0.25 mm ID, and guard column (10 m) (Restek). The initial temperature of GC oven was 60 °C for 60 seconds and the temperature increased to 325 °C (@10 °C/min). Metabolites were identified using NIST library with Xcalibur software.

### Statistical analysis

Statistical analyses used MATLAB, GraphPad Prism (version 10.2) or the R statistics package. Unless otherwise noted, all analyses involved either a two-way full factorial analysis of variance (ANOVA) without or with repeated measures, depending on the specific test (critical α<0.05). Post hoc tests used Tukey’s honest significant difference (HSD) procedure. Where appropriate, false discovery rate (FDR, q ≤ 0.05) correction was used. The resultant statistical test parameters are indicated in figure legends or in the results section. Initial analyses revealed only trends for sex. Male and female data were therefore pooled to increase statistical power. The dependent variables in locomotor tests were sequential and repetitive beam breaks, total, margin, and center distances. Dependent variables for sucrose preference tests are fluid intake consumed in 24 h (ml) and sucrose preference index. For social interaction, the dependent variables are number of ROI entrances, average speed in ROI (in pixels per frame, or ppf), time spent in ROI, and time of initial approach towards ROI. For cued fear conditioning, the dependent variable is freezing counts summed over four 60 second trial bins per day.

Global network measures (i.e., SW, modularity, CPL, efficiency, transitivity, assortativity) were analyzed first as AUC values. Measures with significant differences were again analyzed across density thresholds to determine the threshold level where significant differences are observed. Upon determining the significance of modularity and assortativity, we analyzed the number of modules per group, number of nodes per module, and the normalized rich club index. All single threshold nodal analyses were carried out for a network density threshold of 16%. These included node degree, node strength, participation coefficient, clustering coefficient, and local efficiency. These were corrected for multiple comparisons across 60 nodes using the Benjamini-Hochberg FDR procedure (q ≤ 0.05).

Additional exploratory analyses were carried out on several variables to assess novel links between network measures and affective behavioral measures. For this, we used 9 variables, including AUC values for SW, modularity, CPL, transitivity, assortativity, freezing counts on days 1-3 and margin distance. This data set provided 9 variables with 30 observations each. Factor analysis was first used to explore associations between variables of most importance in the data set (i.e., evaluate variable loadings for the first two factors contributing most to the variance in the data). Projections of variables and data points onto the first 2 dimensions was carried out with principal components analysis (PCA) and cluster analysis using FactoMineR (Lê et al., 2008). A multivariate ANOVA (MANOVA) was used to examine overall differences and report specific univariate associations of potential future interest (α<0.05).

Further analysis of fMRI scans was carried out using probabilistic ICA (Beckmann and Smith, 2004) as implemented in MELODIC Version 3.15, part of FSL. The following data pre-processing was applied to the input data: masking of non-brain voxels; voxel-wise de-meaning of the data; normalization of the voxel-wise variance. Pre-processed data were whitened and projected into a 20-dimensional subspace using PCA. The whitened observations were decomposed into sets of vectors which describe signal variation across the temporal domain (time-courses), the session/subject domain and across the spatial domain (maps) by optimizing for non-Gaussian spatial source distributions using a fixed-point iteration technique (Hyvarinen, 1999). Estimated component maps were divided by the standard deviation of the residual noise and thresholded (0.6) by fitting a mixture model to the histogram of intensity values (Beckmann and Smith, 2004). The resulting components were overlaid on the multi-subject template and classified according to peak z score anatomical location. A series of two-stage multiple linear regressions were used to back-project the group ICA components to subject-specific time courses (using spatial regressions) and spatial components (using temporal regressions). The subject-specific spatial component maps were then used in statistical comparisons for each component in the FSL randomise tool. The statistical design matrix was generated using the FSL Glm tool. We used a one-way ANOVA general linear model design. Randomization tests were carried out with 500 permutations (corrected p-level for significance is 0.05 and statistical thresholding of maps by threshold free cluster enhancement).

For metabolomics data, the intensity of metabolites was imported to the web-based MetaboAnalyst 5.0 platform (https://www.metaboanalyst.ca/) and normalized by the sum of intensities followed by generalized log10 transformation and pareto scaling before multivariate analysis. Further, principal component analysis (PCA), partial least square discriminant analysis (PLS-DA), and Hierarchical Clustering analysis were employed on the dataset (Pang et al., 2021). The Q^2^ and R^2^ values were calculated to assess supervised PLS-DA model. The Variable Importance in Projection (VIP) derived from PLS-DA was used to extract the importance of metabolites to PLS-DA model. Metabolites with a VIP score>1 is weighed to be influential in PLS-DA.

## Results

### 10 wo mice display higher locomotor activity than 30 and 60wo mice

Mice were tested in a novel open field environment one week after arrival. The ages at the time of testing were 11, 31, and 61wo. We used a one-way ANOVA with Tukey’s multiple comparison post hoc test to analyze the data. The 10wo group showed a significantly greater total distance (F_2,27_=4.5 p=0.02; Tukey’s post hoc p=0.03 and 0.04 comparing 10wo vs 30-60wo, respectively) and margin distance than 30 and 60wo mice (F_2,27_=6.3, p=0.006; Tukey’s post hoc p=0.01 comparing 10wo vs both 30-60wo mice) (**Figure 3**). The greater margin distance activity in 10wo mice was driven by female mice (sex main effect F_1,24_=5.9, p=0.02, with margin distance higher in 10wo female mice compared to 30-60wo females and 10wo males; Tukey’s post hoc p <0.05) (**Supplemental Figure 4)**. No significant age or sex differences were observed in other locomotor activity measures, such as sequential and repetitive beam breaks, total and center distance.

**Figure 3.**
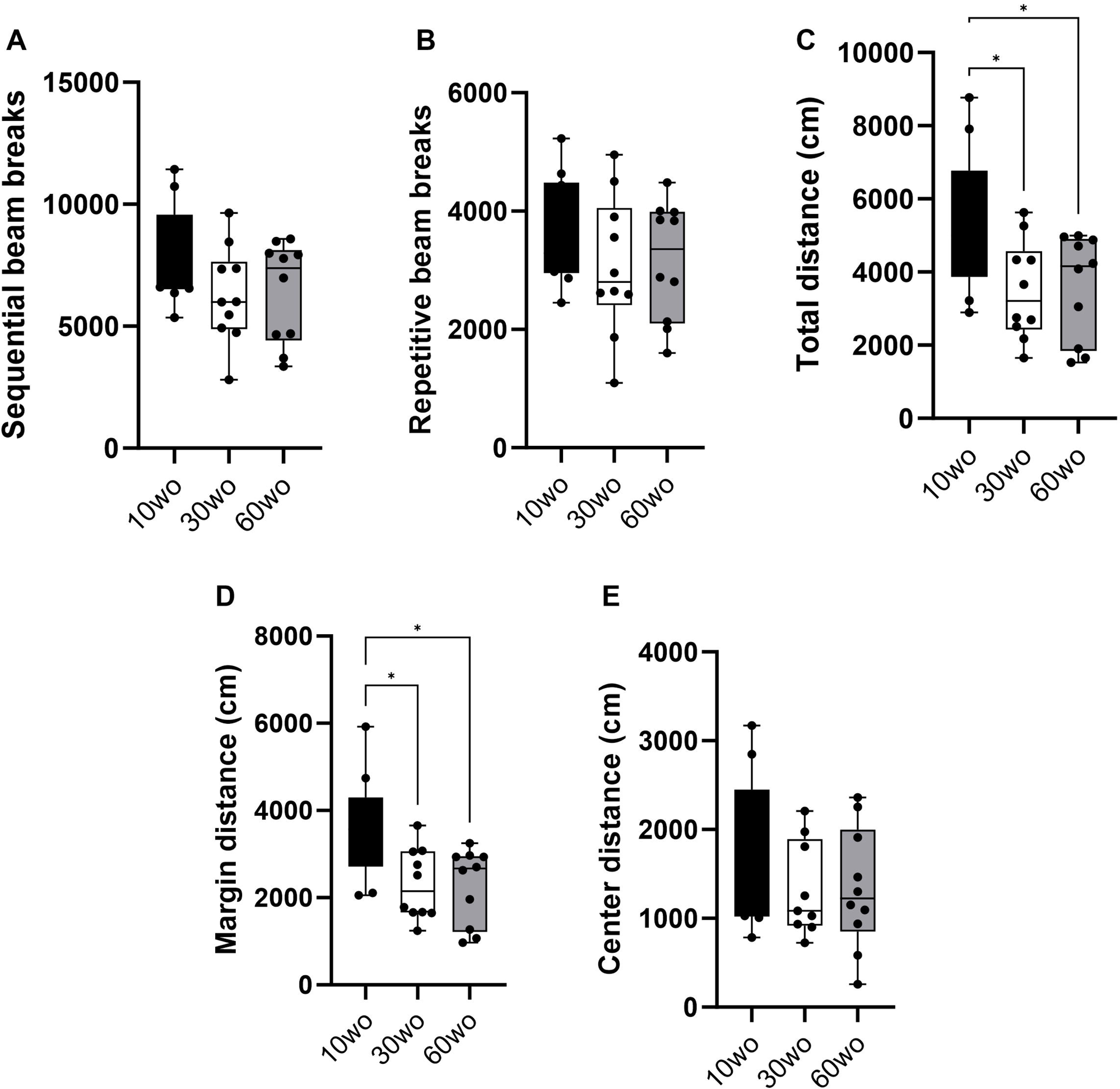
Young 10wo mice showed greater activity in a novel test environment compared to 30 and 60wo mice. A-B) No differences were observed for sequential and repetitive beam breaks activities between the age groups. C) Total distance (in cm) differed between the age groups, with 10wo mice showing a non-significant trend compared to 60wo mice (F_2,27_=4.5 p=0.02; *Tukey’s post hoc p=0.03 and 0.04 comparing 10wo vs 30-60wo, respectively). D) Movement along the margins of the novel test cages different between the age groups, with 10wo mice showing greater movement relative to both 30 and 60wo mice (F_2,27_=6.3, p=0.006; *Tukey’s post hoc p=0.01 comparing 10wo vs both 30-60wo mice). The greater margin activity in 10wo mice was largely driven by female mice (sex main effect F_1,24_=5.9, p=0.02, with margin activity levels higher in female vs male 10wo and 30wo mice, p = 0.004 for female vs male 10 and 30wo mice; n = 10/pooled groups). Data in are box-whisker with interquartile range and minimum-maximum values around the median (dots are data points).

### 10wo mice display brief and faster patterns of social exploratory activity compared to 30-60 wo mice

Sociability tests were carried out when mice were 12, 32, and 62wo. We used a two-way ANOVA (age x test session) with repeated measures. No effect of sex was observed. Thus, data were pooled in our analyses. No age differences in the number of ROI entrances was observed but a significant difference across test sessions was observed (main effect of session F_2,46_=49.4 p<0.00001) (**Figure 4A**). Thus, all age groups entered the ROI where the novel test mouse was located a similar number of times and displayed reductions in the number of entrances during a second encounter with the same test mouse by the same degree. 10wo mice showed a greater mean speed within the ROI than 60wo mice, during the re-encounter with the familiar mouse and exploration of the 2^nd^ novel mouse (F_2,23_=5.9 p = 0.008; Tukey’s post hoc p = 0.02 and 0.03, respectively) (**Figure 4B**). We also observed significant effects of test session and age, but not age x session interactions, on time spent in the test mouse ROI (session F_2,46_=5.5 p = 0.007; age F_2,23_=4.5 p = 0.02) (**Figure 4C**). 30wo mice spent more time in the ROI during the first novel mouse encounter than 10wo mice (Tukey’s post hoc p = 0.007). Only 30wo mice showed reductions in their time spent in the ROI on the second encounter with the familiar mouse and with the new novel mouse (Tukey’s post hoc p = 0.02 and 0.005, respectively). No effects of age or session were observed on time in initial approach (**Figure 4C**).

**Figure 4.**
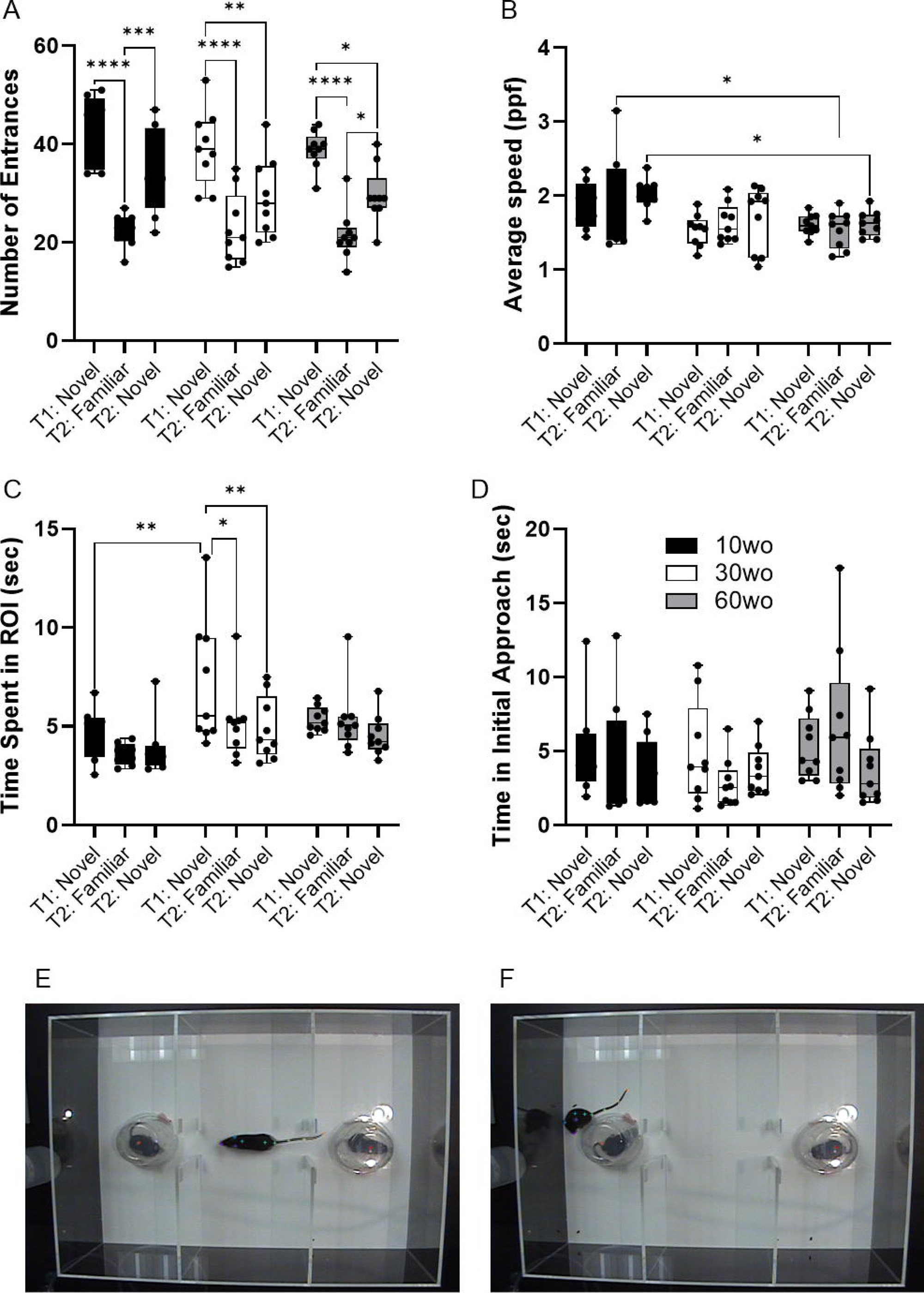
Age differences in sociability towards novel mouse. A) All mice showed similar number of entrances in the novel mouse ROI in the first test (T1), reduced entrances during second exposure to the familiar mouse in a second test (T2) and showed high levels of entrances in a second novel mouse ROI during the second test (T2) (main effect of session F_2,46_=49.4 p<0.00001). Guided asterisks indicate significant differences at *p<0.05, ***p<0.001, and ****p<0.0001 over T1-T2 sessions. B) 10wo mice showed a greater mean speed within the ROI than 60wo mice, during the re-encounter with the familiar mouse and exploration of the 2^nd^ novel mouse (F_2,23_=5.9 p = 0.008; Tukey’s post hoc p = 0.02 and 0.03, respectively). C) The 10wo mice spent less time in ROI where ‘bait’ mouse was located compared to 30wo mice (age F_2,23_=4.5 p = 0.02; Tukey’s post hoc *p<0.05, ***p<0.001). D) No significant differences in time to initial approach. Group sizes are: 10wo n = 8, 30wo n = 9, 60wo n = 9. E-F) Representative video frames capturing exploratory activity by the experimental mouse and novel and familiar mice under inverted clear plastic ‘bait’ cups.

### No age differences in sucrose consumption

Sucrose preference testing was conducted a week after sociability tests when mice were 13, 33, and 63wo. No effect of sex was observed. Thus, data were pooled in our analyses. All mice consumed more sucrose solution than water. No age-related differences in the amount of sucrose and water consumed nor differences in the sucrose preference index were found (**Supplemental Figure 5**). We also analyzed sucrose consumption during the 3 days of acclimation and observed no differences between the groups (**Supplemental Figure 5**).

### 60wo adult mice display high variability in their freezing responses during initial fear conditioning but express fear memory at 24 and 48 h

Fear conditioning was conducted a week after completing fMRI scans when mice were 16, 36, and 66wo. Percent time freezing was analyzed using a two-way repeated measures ANOVA. No effect of sex was observed. Thus, data were pooled in our analyses. On day 1 of fear conditioning, all age groups progressively developed unconditioned-conditioned stimulus (US+/CS+) associations (**Figure 5**). We observed a rise in percent time freezing over the course of each tone-shock exposure in all age groups (effect of trial F_3,81_ = 66 p < 0.0001 and age F_2,27_ = 7.5 p = 0.0025; **Figure 5A**). 60wo mice showed higher percent freezing across cue-shock trials than the other age groups (Tukey’s post hoc test indicated p<0.05 comparing 60wo to 10wo and 30wo mice on trials 1-3 and p = 0.02 compared to 10wo mice on trial 4), although the freezing responses were more variable in the 60wo mice than in the other two age groups. Data were next analyzed as average percent time freezing per day (**Figure 5B**). When compared across days, all age groups showed robust recall of fear conditioning, observed as high levels of freezing in response to the tone alone, at 24 and 48 h (main effect of age F_2,27_ = 4.6 p =0.02; main effect test day F_2,54_ = 66.8 p < 0.0001). At 24 h, all mice maintained a high percent freezing relative to day 1 (Dunnett’s test p <0.05 in all age groups comparing day 2 to 1). At 48 h, the cages were modified such that the tone was delivered in a context with different sensory features (walls and floors were smooth white plastic in a triangular configuration, as reported in (Anagnostaras et al., 2010). Still, 10wo and 60wo mice maintained high levels of freezing in response to the conditioned cue, relative to day 1 (Dunnett’s test p < 0.05). The 30wo mice showed a similar but non-significant trend towards increased fear recall at 48 h (**Figure 5B**). The freezing responses by 60wo mice were higher than those of 10wo mice on the 3^rd^ day of testing (Dunnett’s test p < 0.05).

**Figure 5.**
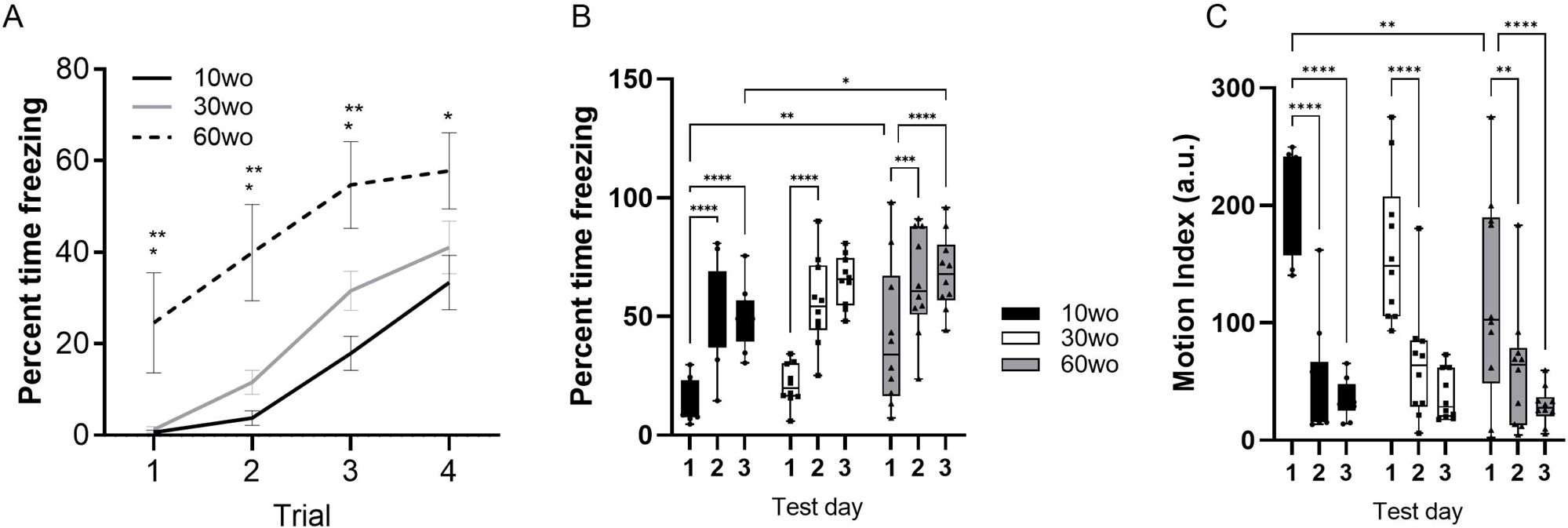
60wo mice demonstrate high freezing responses at 24 and 48 h post despite variable initial development of conditioned freezing responses. A) Although all mice develop conditioned freezing by trials 3 and 4, the oldest age group showed high levels of freezing compared to 10-30wo mice (Tukey’s HSD p<0.05 comparing 60wo to 10* and 30wo** mice on trials 1 and 2). B) Average percent time spent freezing over 3 days of testing (conditioning, recall at 24h, recall in modified context at 48h). All mice show high levels of freezing at 24 and 48 h, although in comparison to 10wo mice, 60wo mice maintain high freezing responses in a modified context on day 3 (asterisks indicate significant differences at *p<0.05, **p<0.01, ***p<0.001, and ****p<0.0001, Tukey’s post hoc test). C) Motion index (arbitrary units, a.u.) over 3 days of testing. All mice show reductions in general locomotor activity inside the operant cages on days 2-3 compared to day 1 (asterisks indicate significant differences at **p<0.01, and ****p<0.0001, Tukey’s post hoc test). Data in A are mean standard ± error and in B-C are box-whisker with interquartile range and minimum-maximum values around the median (dots are data points; n=10/group).

We analyzed a motion index as a surrogate measure of general locomotor activity using an infrared motion sensitive camera inside the test cages. Consistent with results in **Figure 3**, 10wo mice showed higher motion index levels than 60wo mice in the novel test cage on day 1 (**Figure 5C**) (day by age interaction F_4,54_ = 2.7 p =0.04, Tukey’s HSD p = 0.008 day 1 comparing 10wo vs the 60wo group; main effect test day F_2,54_ = 84.9 p < 0.0001). The motion index was significantly reduced in all groups by days 2 and 3 of testing (Tukey’s HSD p < 0.05 comparing days 1 vs 2 and 3 for all age groups). A closer assessment of the motion index over the tone-shock presentation trials on day 1 and tone responses on day 2 provided further evidence of highly variable responses to conditioned and unconditioned stimuli in 60wo mice compared to the other two age groups (**Supplemental Figures 6-8**). The 60wo mice did have a reduced initial motion response to the tone, but this was only significant during the second tone-shock presentation (two-way ANOVA with repeated measures showed effect of time, F_2.8,76.1_=46 p<0.0001, and age, F_2,27_=7.5 p = 0.0025, but no interactions; Tukey’s multiple comparison test indicated p<0.05 comparing 60wo to 30wo, on tone-shock trial 2). However, all mice responded to tone, shock, and showed reduced motion during baseline over the course of all 4 trials on day 1 (**Supplemental Figures 6D and 8**). Further declines in motion were not observed on day 2 (**Supplemental Figure 7D**), suggesting acclimation of motion overlapping with conditioned fear learning.

### Young adult mice show a higher network modularity compared to older adult mice

Functional MRI scanning was carried out over a two-week period following sucrose preference testing. Mice were 14-15, 34-35, and 64-65wo at the time of scanning. Data were analyzed using two-way ANOVA with repeated measures and Tukey’s multiple comparison test. No effect of sex was observed. Thus, data were pooled in our analyses. Analysis of network measures obtained from functional connectivity matrices indicated significant main effects of age, but not sex, for SWC (age by threshold interaction F_26,312_ = 2.4 p = 0.0002), modularity (age by threshold interaction F_26,312_ = 1.79 p = 0.01), and assortativity (age by threshold interaction F_26,312_ = 2.6 p = 0.00006) (**Figure 6**). Tukey’s post hoc comparisons further revealed that SWC was highest in 30wo mice relative to 10wo mice and both modularity and assortativity were highest in 10wo mice relative to 30 and 60wo mice (**Figure 6**). Significant pairwise differences were observed across several edge density thresholds (see **Figure 6C-6E**; details in figure legend). We analyzed the number of modules (at 16% edge density) per age group and observed a significantly greater number of modules in 30 vs 10wo mice (**Figure 6A** and **6F**) (Kruskall-Wallis test p = 0.02). Network randomization removed these differences (**Figure 6B**). The number of nodes per module did not differ between groups (**Figure 6G**). To confirm the significant age related changes in functional network modularity, we further investigated 5 additional community detection routines available in igraph, including greedy optimization (Clauset et al., 2004), edge betweenness (Newman and Girvan, 2004), random walk (Pons and Latapy, 2005), lead Eigenvector (Newman, 2006), and a different implementation of Louvain’s algorithm (Blondel et al., 2008) (**Table 1**). These were all estimated at 16% edge density. Across most of these algorithms, the number of modules detected was lowest in 10wo mice compared to the other two age groups. In addition, the modularity index was highest in 10wo mice compared to the two older age groups (**Table 1** provides statistical results).

**Figure 6.**
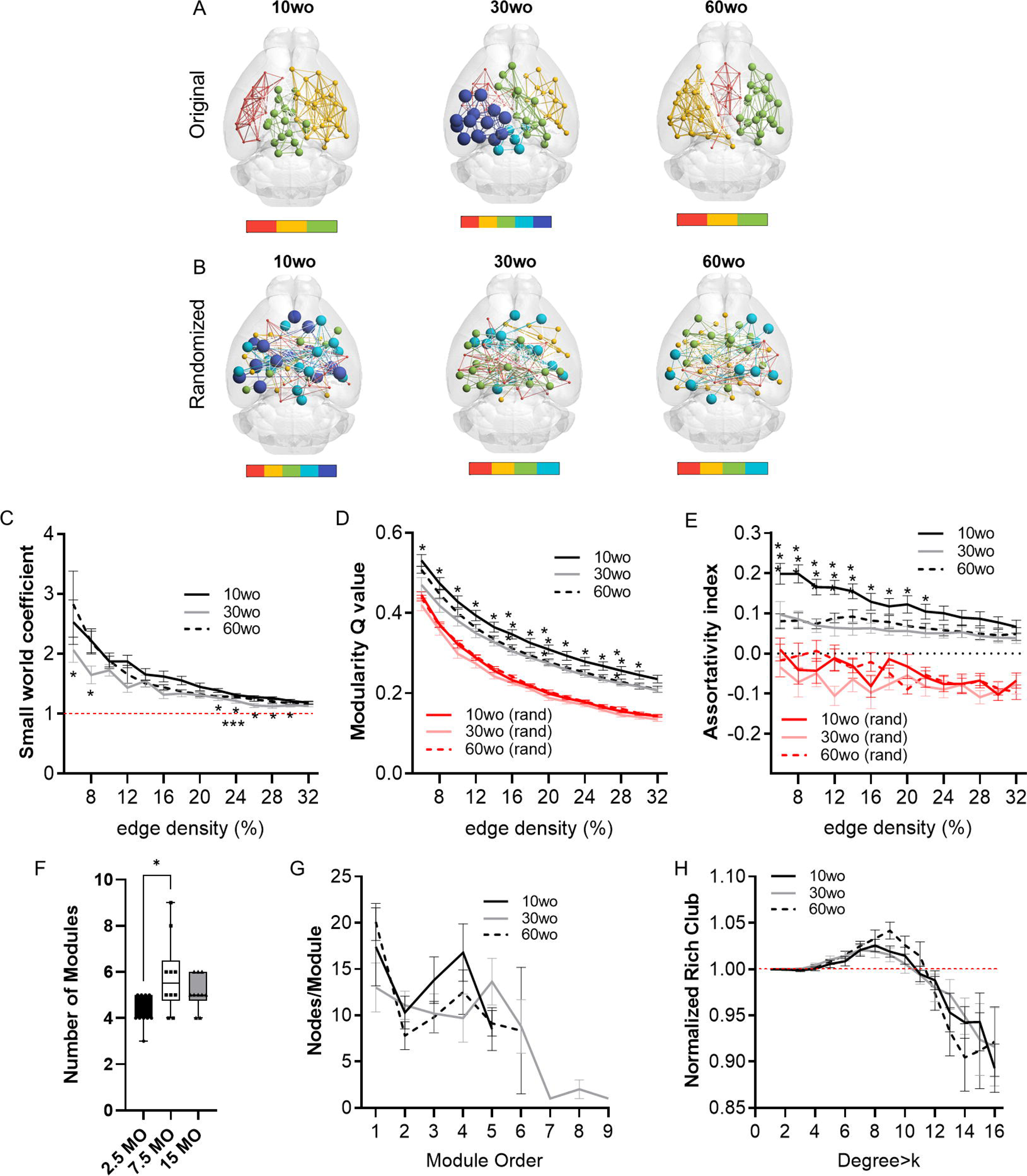
Young 10wo mice had greater modularity and assortativity of functional connectivity networks than 30 and 60wo mice. A) 3D functional connectome maps illustrating mean modularity value (node size) and node community affiliation (node and edge color) for 10, 30, and 60wo mice (density threshold = 16% edges retained). The bar below maps indicates the number of identified modules. B) Randomized 3D connectome maps for the same data in A. C) Small world coefficient was lowest in 10wo mice (age by threshold interaction F_26,312_ = 2.4 p = 0.0002, Tukey’s HSD * and *** p<0.05 comparing 10wo and 30wo to 60 weeks, respectively). D) Modularity was highest in 10wo mice (age by threshold interaction F_26,312_ = 1.79 p = 0.01, Tukey’s HSD * and ** p<0.05 comparing 10wo to 30wo and 60 weeks, respectively). E) Assortativity was highest in 10wo mice (age by threshold interaction F_26,312_ = 2.6 p = 0.00006, Tukey’s HSD * and ** p<0.05 comparing 10wo to 30wo and 60 weeks, respectively). F) Number of modules identified at density-16% was higher in 30wo compared to 10wo mice (Kruskall-Wallis test p = 0.02). G) No difference in number of nodes per module was observed. H) No difference in the normalized rich club coefficient was observed. N = 10/group. Data in A-E and G-H are mean standard ± error and in F are box-whisker with interquartile range and minimum-maximum values around the median (dots are data points).

At 16% edge density, we also observed significant main effects of age on transitivity and assortativity (**Table 1**). Transitivity was lowest in 60wo vs 30 and 10 wo mice. assortativity was significantly greater in 10wo vs 30wo mice. This was observed using Brain Connectivity Toolbox’s routine for calculating network assortativity. Although a similar trend was observed using igraph, it did not reach statistical significance (**Table 1**).

### Young adult mice have robust functional connectivity between affective brain regions in auditory/temporal cortical and striatal networks

Group ICA identified 9 of 20 functional network components similar to previously identified in mice (Jonckers et al., 2011;Grandjean et al., 2014a;Sforazzini et al., 2014;Bukhari et al., 2017). The ICA components included anterior and posterior cingulate, somatosensory, visual, auditory/temporal, piriform/amygdala, hypothalamic and dorsal striatal networks (**Supplemental Figure 9**). **Figure 7** shows the results for statistical comparisons between age groups. The 10wo mice had greater functional connectivity in the auditory/temporal network than 60wo mice. The areas showing greater connectivity within the right hemisphere of this network in 10wo mice included anterior cingulate, auditory/temporal subregions, ventral hippocampus, dorsal striatum, midbrain reticular nucleus, bed nucleus of stria terminalis, posterior thalamic areas, and superior colliculus. The left hemisphere had less differences and included primary and secondary somatosensory areas, striatum, visual cortex, hippocampus, and thalamic subregions (**Figure 7**). The left hemisphere striatal network had greater functional connectivity in 10wo compared to 60wo mice. The greater functional connectivity in the left striatal network of 10wo mice included piriform, hippocampus and superior colliculus. We also observed greater functional connectivity in a left hemisphere striatal network in 30wo compared to 60wo mice and in a right hemisphere somatosensory network in 10wo compared to 30wo mice.

**Figure 7.**
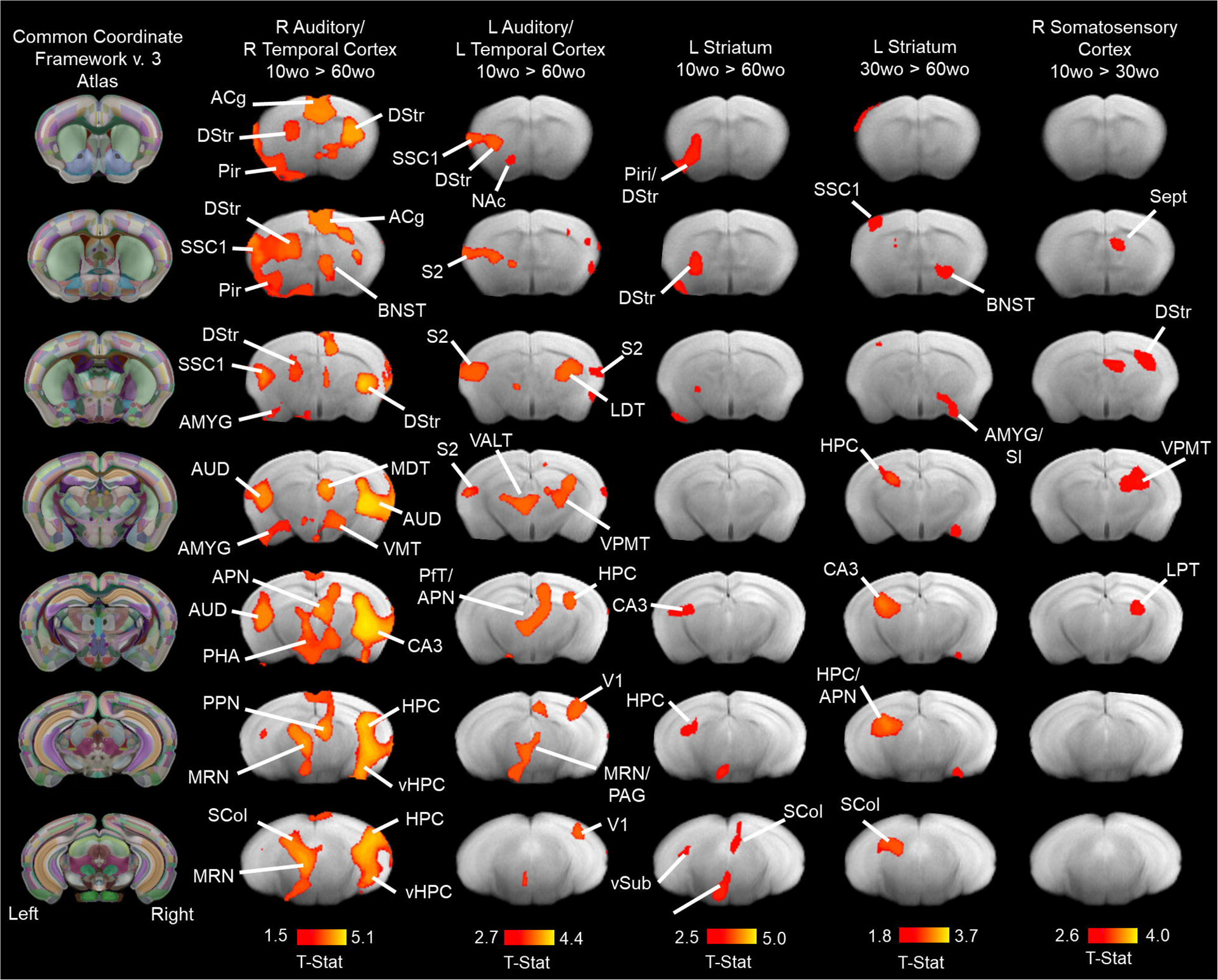
Age associated reductions in functional connectivity in primary sensory and association cortical areas and striatum. Lines indicate regions of the brain with statistically significant differences in connectivity with the network component (indicated above maps). Statistical analysis used a one-way ANOVA with mixed effects (p<0.05, cluster corrected; higher level permutation tests with 5000 permutations; n = 10/group). Maps thresholded at t >2.3 (bar graph below maps) and overlaid onto multi-subject T2 anatomical template. Titles over the maps indicate the original network component identified by group pICA (see methods for details). Column maps on the left correspond to CCFv3 atlas of the mouse brain. Abbreviations: ACg, anterior cingulate; AMYG, amygdala; APN, anteroposterior nucleus of thalamus; AUD, auditory cortex; BNST, bed nucleus of stria terminalis; CA3, ‘Ammons Cornus’ 3 subfield of hippocampus; DStr, dorsal striatum; HPC, hippocampus; LDT, laterodorsal thalamus; MDT, mediodorsal thalamus; MRN, median Raphe nucleus; NAc, nucleus accumbens; PHA, posterior hypothalamic area; Pir, piriform cortex; PfT, parafascicular thalamus; S2, secondary somatosensory cortex; Sept, septum SCol, superior colliculus; SI, substantia innominate; SSC1, primary somatosensory cortex; V1, primary visual cortex; VALT, ventral anterolateral thalamus; vHPC, ventral hippocampus; VMT, ventromedial thalamus; VPMT, ventral posteromedial thalamus; vSub, ventral subiculum.

### Statistical interactions between age, locomotor activity, fear conditioning and indices of functional network connectivity

MANOVA revealed a significant effect of age across several connectomic and behavior variables (F_9,20_=4.6, p=0.002). Principal components analysis (PCA) revealed a relatively strong relationship between several of these variables. This is observed by contributions to the first 2 dimensions, which jointly explain 56.08% of variance in the data. To establish significance, PCA results for the first 2 dimensions (56.08%) were compared against a reference value (43.65%) representing 95% quartiles of distribution of percent explained variances (inertia) of simulated data tables of equivalent size and normal distribution (**Figure 8**). The positive direction along dimension 1 opposes 90% of mice in the youngest group (1-8 and 10 are in the youngest 10mo group) from many of the mice in the other two age groups (**Figure 8A and Supplemental Figure 10A**). This positive direction is characterized by high values freezing counts on days 2-3 of fear conditioning and high margin distance activity in a novel open field environment and high values for small world coefficient, transitivity, and modularity in 70% of the 10mo mice (**Supplemental Figure 10B**). Cluster dendrogram in **Figure 8B** classified 3 clusters. A majority of young 10mo mice were classified in cluster 3 (green). This group is again characterized by high values for modularity, transitivity, small world coefficient, margin distance activity and days 2-3 of fear conditioning.

**Figure 8.**
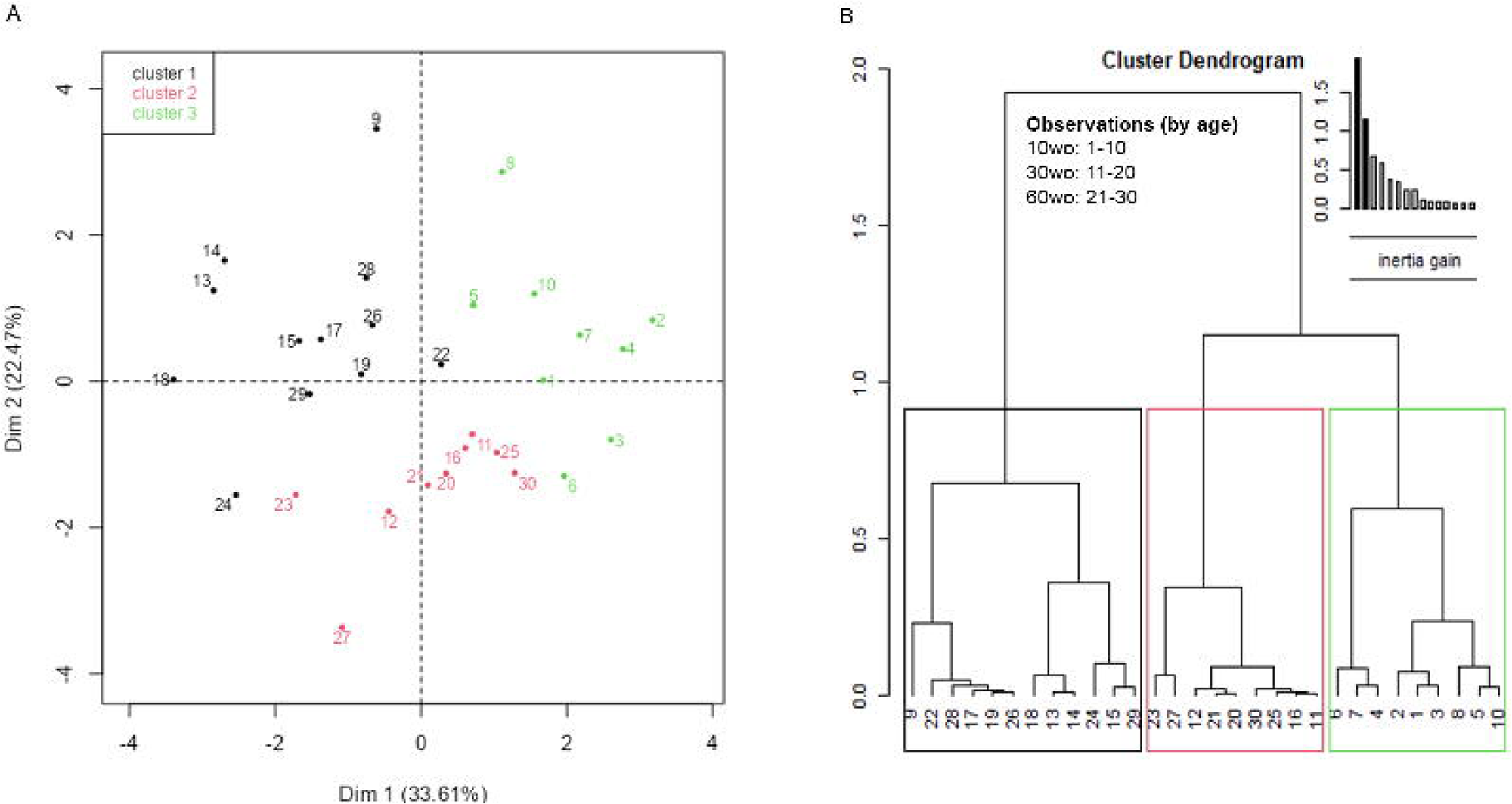
Exploratory multivariate analyses reveal significant associations of interest between functional network measures and affective behavior variables. The first two dimensions of the PCA explain 56.08%, which is above 95%-quartiles of random distributions predicting 43.62% of variance explain with equal size data tables. A) Individual factor map with labeled individuals contributing to a high degree to the first two dimensions. B) Cluster tree partitioning data by ascending hierarchical classification of principal components. Cluster 3, made mostly of individuals in the youngest age group, is characterized by high values for modularity, transitivity, small worldness, marginal distance activity and fear recall (**Supplemental Figure 10**).

### Metabolomic profiles in prefrontal cortex and hippocampus differ between 10, 30 and 60wo mice

Results for PCA and partial least squares discriminant analysis (PLS-DA) for GC-MS data obtained from the mouse hippocampus are shown in **Supplemental Figure 11** and **Figure 9** (panels A and B). Variable importance in projection (VIP) scores plot of the PLS-DA model for mouse hippocampus indicated differences in the top 25 metabolites between 2.5, 7.5, and 15 mo mice. Among the top 25 listed in **Figure 9** (panel A), methionine and cysteine had the highest VIP scores. The top 7 had a bidirectional transition in relative levels when youngest mice having lowest or highest levels and late middle-aged mice having diametrically distinct levels. This relationship included methionine, cysteine, malate, fumarate, succinate, N-acetylglycine, alanine, and taurine. The differential prominence of these metabolites across age groups is supported by clustering in heatmap shown in **Figure 9** (panel B). Results for PCA and PLS-DA for GC-MS data obtained from the mouse midline prefrontal cortex are shown in **Supplemental Figure 12** and **Figure 9** (panels C and D). VIP scores plot of the PLS-DA model for mouse prefrontal indicated differences in the top 25 metabolites between 2.5, 7.5, and 15 mo mice. Among the top 25 listed in **Figure 9** (panel C), proline, taurine, and methionine had the highest VIP scores. The differential prominence of these metabolites across age groups is supported by clustering in the heatmap in **Figure 9** (panel D).

**Figure 9.**
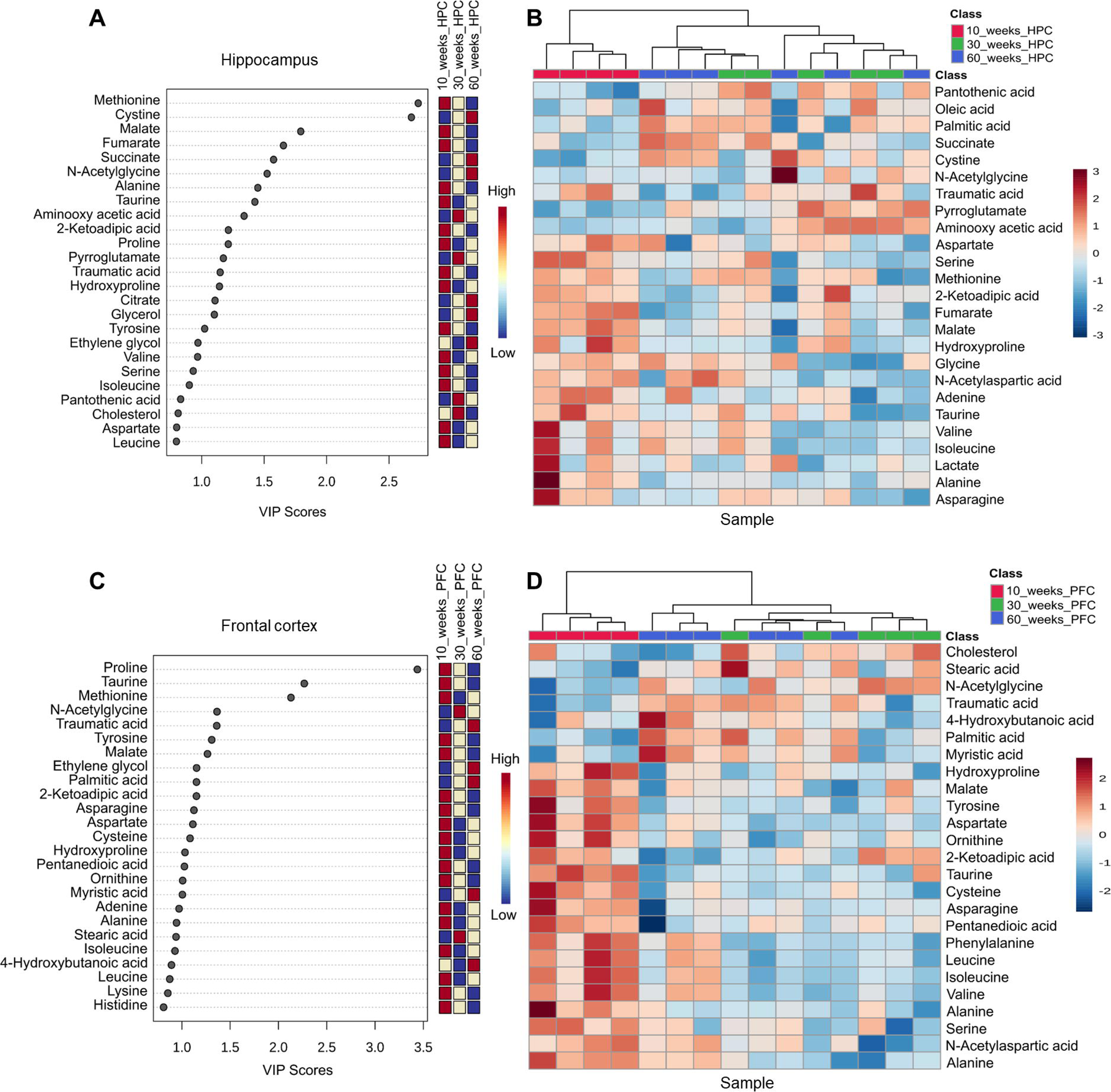
Mouse brain metabolomics identifies age variations in methionine, cysteine, and proline in prefrontal cortical and hippocampal regions. A-B) Variable importance in projection (VIP) scores plot of the PLS-DA model for mouse hippocampus indicated differences in the top 25 metabolites between 10, 30, and 60wo mice). The differential prominence of these metabolites across age groups is supported by clustering in heatmap. C-D) VIP scores plot of the PLS-DA model for mouse prefrontal indicated differences in the top 25 metabolites between 10, 30, and 60wo mice. D) The differential prominence of these metabolites across age groups is supported by clustering in the heatmap. Group sizes are: 10wo n = 4, 30wo n = 5, 60wo n = 6.

## Discussion

Our present results indicate that 10wo, 30wo and 60wo C57BL/6J mice differed in their locomotor activity in a novel environment, social interaction time and speed, and fear conditioning. These differences were largely observed between the youngest and oldest age groups and may reflect differences in ‘emotionality’ between the two age groups. The youngest mice appear more active in response to novelty, such as a novel environment or a novel social interaction. The oldest group appeared less active and exhibited a more ‘cautious’ and variable pattern of behavior during fear conditioned training. Our fMRI results indicate that young 10 wo mice had greater functional connectivity between a temporal cortical/auditory cortex network and subregions of the anterior cingulate cortex and ventral hippocampus, among other regions, and a greater network modularity and assortative mixing of nodes compared to older adult mice. We observed a close correspondence between behavioral measures of fear conditioning, locomotor activity in a novel environment and measures of network efficiency and segregation in young mice, which was no longer apparent in older adult mice. Finally, metabolome analyses revealed significantly different neurochemical profiles in prefrontal cortex and hippocampus between 10wo mice and the other age groups, with several essential amino acids differing the most between the groups (e.g., methionine, cysteine, proline, and taurine).

In mice, conditioned fear memory varies with age in both context (hippocampal-dependent) and cued (amygdala-dependent) paradigms. Compared to adult 6mo mice, late middle age 17mo C57BL/6 mice show deficits in renewal of extinguished freezing in a modified context (Sanders, 2011). Eighteen-month-old mice showed greater freezing in response to context but lower freezing in response to tone than 3mo mice (Evans et al., 2021). These results suggest dissociable aging changes in conditioned fear memory, perhaps involving distinct neural circuits involving amygdala and hippocampus (Evans et al., 2021). Twenty three month old mice showed higher cued fear compared to 2mo mice but impaired renewal of conditioned freezing following extinction than 12mo mice (Oh et al., 2018). This was associated with altered stress reactivity (Oh et al., 2018). Using a trace eye blinking conditioning protocol, Kishimoto showed significant impairments in conditioned responses in aged 20mo compared to 12 and 2mo mice (Kishimoto et al., 2001). The above studies support differences in conditioned fear in aged versus young mice. Age differences in hippocampal/prefrontal structure and function at the neuronal and synaptic level, stress coping/stress molecular signaling changes and changes in neuroimmune activity are thought to account for the observed age differences in conditioned freezing in mice (von Bohlen und Halbach et al., 2006;Oh et al., 2018;Evans et al., 2021). Viewed from this perspective, human studies reporting lower amygdala or cingulate activation in response to negative valence stimuli in aged versus young adults may be accounted for by deficits in these brain areas as well (as opposed to a greater reactivity in young adults) (Chaudhary et al., 2023). Our present data reveals a close correspondence between conditioned fear recall and modularity, transitivity, and small world coefficient (**Figure 8** and **Supplemental Figure 10**). It should be noted that there is extensive literature on the role of the amygdala and its subregions in fear conditioning. One of the limitations of this study is the use of an fMRI sequence based on the gradient recalled echo that, while having signal to noise benefits, leads to signal loss in air-tissue interface regions where the amygdala is located. This effect can be appreciated from the representative cropped fMRI scan in **Figure 2**. Therefore, the lack of results in the amygdala should therefore be interpreted with caution.

Young 10wo mice displayed greater locomotor activity than 30 and 60wo mice, consistent with evidence of decreasing locomotion with age (Elias et al., 1975). Male B6 mice are generally more active at 1-3mo than at 7 or 13mo (Lhotellier and Cohen-Salmon, 1989;Serradj and Jamon, 2007), although others have reported greater locomotor activity in 17mo vs 5 and 25 mo B6NIA mice (Frick et al., 2000), greater locomotor activity in 3-8mo versus 28 mo B6NJ (Fahlstrom et al., 2011), or no difference between 2 and 18mo B6 mice (Barreto et al., 2010). In our present observations, locomotor activity in the younger B6 group was largely along the margins of a novel test cage. This suggests novelty associated exploratory activity along the safer walls of the cages in 10wo mice. Under novel exploratory conditions, however, 24mo B6 mice were observed to display greater exploratory activity than 2mo mice (Ammassari-Teule et al., 1994). Open field activity in our present work was greatest in 10mo mice and specifically female mice. Thus, sex differences also should be considered when assessing affective behaviors (Knight et al., 2021).

Our results showed no differences in sucrose preference between the studied adult age groups. All mice consumed preferred sucrose solution over water to the same degree. This result contrasts with previous reports. Eighteen-month-old (∼72wo) C57BL/6N mice reportedly show less sucrose reward preference than 3mo mice (Malatynska et al., 2012). This was correlated with other signs of depressive-like behaviors in aged versus young mice (Malatynska et al., 2012). Similar reductions in sucrose and milk reward preference in 13-15mo versus 3mo C57BL/6 were reported (Harb et al., 2014). Conversely, aged 24mo Balb/c mice did not show differences in sucrose preference, locomotor activity, immobility in a tail suspension test compared to 4-6mo mice (Kelley et al., 2013). Discrepancies between studies may arise from differences in the sucrose preference protocol, differences in mouse strains, and age groups. We also examined social interactions in mice. While all mice showed similar social interactions in terms of the number of approaches towards test mice, 10 wo mice display faster and more brief interactions compared to the older mice. This is partly consistent with results from Shoji and Miyakawa, who reported similar levels of social interactions in B6 mice ages 2-23mo. However, they reported reduced preference for novel social interactions in 17 and 23mo relative to 11mo old mice. Although we observed no such preferences using a supervised learning approach to assess social interactions, 60wo mice did display slightly lower novelty preferences during a second novel social encounter relative to a first novel encounter (**Figure 4**).

We observed differences in functional connectivity within a temporal/auditory cortex network. Specifically, anterior cingulate cortex and ventral hippocampal connectivity within this network was strongest in 10wo compared to 60wo mice. Our results using functional MRI agree with previous structural neuroimaging of mice along the same age range, which identified maturational changes from young to middle age in auditory, cingulate, hippocampal formation, and somatosensory (parietal) cortices (Clifford et al., 2023). The results in our present study showing stronger temporal/auditory connectivity with the ACg and vHPC in the young age group is consistent with the robust cued fear conditioning these mice developed compared to older age groups. These structures have been studied for their roles in generalized fear and anxiety in murine models (Ortiz et al., 2019). Increased synaptic activity in projections from ACg to vHPC regulate the long-term generalization of fear memories (Bian et al., 2019). The vHPC in mice contains neurons that show higher levels of neuronal activity and conditioned freezing behavior in response to threat cues than safety or combined threat/safety cues (compound cues) (Meyer et al., 2019). Conversely, a subset of prelimbic cortical projecting vHPC neurons show increased neuronal activity with safety cues and this was accompanied by lower levels of freezing relative to their response to threat cues (Meyer et al., 2019). Thus, the stronger functional connectivity observed in the present study in young adult mice may be associated with the fear and anxiety related functions of this vHPC-PFC circuit. Importantly, we observed the higher functional connectivity between temporal/auditory and vHPC-ACg regions in young versus older mice prior to their exposure to fear conditioning, thus averting effects introduced by learned fear on functional connectivity. We should point out that the age differences in functional connectivity between ACg, vHPC and other regions within the temporal/auditory network (**Figure 7**) were lateralized, with a much greater functional connectivity observed in the right versus left temporal cortical network. Fear-related anterior cingulate activity has been previously reported to be lateralized in mice (Kim et al., 2012). Inhibition of the right but not left ACg resulted in reduced observational fear learning relative to control mice (Kim et al., 2012).

We also observed a greater functional connectivity within a left dorsal striatal network in 10wo and 30wo compared to 60wo mice. The regions observed to show significantly greater functional connectivity with the dorsal striatum included a subregion of the HPC, superior colliculus, bed nucleus of stria terminalis. The role of the dorsal striatum in regards to fear and anxiety are supported by recent work indicating its synaptic connectivity arising from apical intercalated cells in the amygdala, which regulate fear conditioning in mice via inputs from thalamic, temporal association cortex and basolateral amygdala nuclei (Asede et al., 2021). Finally, we should note that age differences between young (1-2mo) and older adult (5-19mo) wildtype littermates of the transgenic Arcβ familial amyloidosis mouse have been reported (Grandjean et al., 2014b). Young adult mice around 2 months of age had weaker functional connectivity within subregions of the parietal, cingulate and dorsal striatal areas than older mice at 5mo and older (Grandjean et al., 2014b). These results identify similar regions to the ones reported here. However, whereas we observed decreases with age, Grandjean and colleagues (Grandjean et al., 2014b) observed increases in control aging mice. While this may arguably be due to differences in anesthesia protocol (isoflurane 1.4% in Grandjean *et al*. and 0.25% isoflurane and dexmedetomidine used here), it is more likely that differences are associated with the background strain of mice. Mouse strain-dependent genetics likely weighs heavily on brain microstructure and function (Neuner et al., 2019;Ashbrook et al., 2021;Murdy et al., 2022).

Our results indicate a higher functional network modularity and assortative mixing of nodes of young versus older adult mice. The modularity changes we observed in young versus older adult mice were of particular interest since it persisted across several different modularity algorithms. Graph analysis of brain structural covariance matrices of 2, 10, and 22mo mice showed reductions in node degree values from young to middle age in ACg, auditory and somatosensory cortices and an increase in hippocampus (Clifford et al., 2023). They also reported reduced modularity from 2 to 10mo (Clifford et al., 2023). In human subjects, young and early middle age groups have higher local nodal efficiency, clustering coefficients, and path lengths than late middle age and elderly groups (Wang et al., 2023). These graph measures, which suggest changes in network efficiency from young adulthood through aging, were considered in the context of cognitive changes and not affective behaviors. In the present work, we observed statistically meaningful associations between the multiple affective behavioral and connectomic variables. Although this does not confirm causation or even correlation between the variables, the results do point to interactions of interest that should be considered with other circuit specific approaches to determine the relationship between network configurations and affective behaviors.

Finally, we observed age related changes in several amino acid substrates in prefrontal and hippocampal tissue, which supports biochemical changes in these regions that might be linked to the observed maturational changes in brain function and behavior. Each of the substrates that differed the most between the groups are known to play important roles in neuronal function and development. Proline is important for glutamate homeostasis following methamphetamine challenge (Jones et al., 2021). Taurine is involved in developmental changes to neurons in the hippocampus, cerebellum, hypothalamus, and can act via GABA_A_ receptors to modulate firing of GABA neurons (Curran and Marczinski, 2017). Methionine is associated with a host of distinct functions related to the control of gene expression in neurons and high methionine diets have been linked to neuronal damage and accelerated signs of senescence in animal models (Pi et al., 2021). The D enantiomer of cysteine is found in neuroprogenitor cells and is critical to early brain development (Semenza et al., 2021). Taken together, these results suggest ongoing maturational developmental changes in prefrontal cortex and hippocampus in young adult mice as well as senescence-related neurochemical changes in late middle-aged mice.

## Acknowledgements

This study was funded by a National Institute on Aging grant (R21AG065819). Support was also provided by the McKnight Brain Institute of the University of Florida. Joy Buraima was supported by the Summer Neuroscience Internship Program (SNIP) of the McKnight Brain Institute. The contents of this manuscript are solely the responsibility of the authors and do not necessarily represent the official views of the funding agencies. This work was performed in the McKnight Brain Institute at the National High Magnetic Field Laboratory’s AMRIS Facility, which is supported by National Science Foundation Cooperative Agreement No. DMR-1644779 and the State of Florida.

## Scope statement

The study reveals age associated differences in locomotor activity in a novel environment, social interactions, and the development of fear condition. Young mice display greater functional connectivity between specific brain regions responsible for emotion processing, the anterior cingulate, hippocampus, and temporal cortex than late middle-aged mice. Metabolome analysis further identified differences in essential amino acids in prefrontal cortex and hippocampus between age groups. These findings shed light on the impact of aging on emotional responses and provide insights into brain activity as the brain matures.

