## Supplemental Figure legends for "Age-Related Differences in Affective Behaviors in Mice: Possible Role of Prefrontal Cortical-Hippocampal Functional Connectivity and Metabolomic Profiles"

**Supplemental Figure 1.** Thresholding graph density at 16% retains significant edge weights (Pearson r correlation coefficients). A) A graph density of 16% (δ=0.16) corresponds to a mean node degree of 9.1, mean node strength >2.4 and <4.6, and a r ~0.15-0.42. B) Thresholding graphs based on statistically significant Pearson r values (within 95% confidence bands) results in r values between 0.06-0.07 (excluding 2 outliers in 7.5 MO group), density between 40%-95%, node degree between 20-60 and node strength between 4-18. Data in are box-whisker with interquartile range and minimum-maximum values around the median (dots are data points).

**Supplemental Figure 2.** Randomization of functional connectivity networks results in differential organization of connectivity between original and random graphs, while preserving node weights, degree, and strength distributions. A-F) Histograms of node strength and degree distributions at 3 density thresholds (δ=0.08, 0.16, 0.32). Relaxation of graph densities (δ=0.32) adds ‘spurious’ correlations shaping ‘random-like’ distributions in histograms. G-H) Node strength and degree across 60 nodes (at δ=0.16) shows preservation of values between original and randomized networks. I) High correlation between input and output strength sequence accuracies with the randomization of signed graphs (i.e., positive, and negative weights). J-K) Node strength values (sphere sizes at each anatomical location) were preserved while organization of edges in 3D maps in J and K differed. See text for further details.

**Supplemental Figure 3.** Schematic of GC-MS workflow.

**Supplemental Figure 4.** Sex differences in locomotor activity in a novel test environment. A-B) No sex differences in sequential and repetitive beam breaks. C) 10 wo female mice showed greater total distance movement than age-matched male mice (Tukey’s post hoc p = 0.04 comparing 10wo females to 10wo males). D) No sex differences in center distance movement. E) 10 wo female mice showed greater margin distance movement than age-matched male mice (Tukey’s post hoc p = 0.01 comparing 10wo female to 10wo males). Data in are box-whisker with interquartile range and minimum-maximum values around the median (dots are data points).

**Supplemental Figure 5.** No difference in reward sensitivity between age groups. A) Sucrose intake from the left-side cage bottle during the 3 days of acclimatization. B) Sucrose intake from the right-side cage bottle during the 3 days of acclimatization. C) No effect of age or sex was observed on amount of water and sucrose consumed in 24 h. A main effect of type of fluid (sucrose vs water) was observed, with all mice consuming more sucrose solution than water (F_1,23_ = 120 p< 0.0001; *p<0.05 Tukey’s HSD). D) No difference in sucrose preference index was observed. Group sizes are: 10wo n = 8, 30wo n = 9, 60wo n = 9. Data in A-B are mean ± standard error and in C-D are box-whisker with interquartile range and minimum-maximum values around the median (dots are data points).

**Supplemental Figure 6.** Averaged raw motion index traces over the course of day 1 of fear conditioning illustrates consistency in response to electrical stimulation. Traces in arbitrary units were exported and normalized to allow qualitative comparisons. Blue arrows indicate the onset of tone and red arrows the onset of the electric grid floor stimulus. A) Motion traces for 10wo mice. B) Motion traces for 30wo mice. C) Motion traces for 60wo mice. Note the the rise in motion activity in response to the tone onset across the 4 trials in 10-30wo but less so for 60wo mice. 60wo mice show more variability, particularly in the first two trials. Data are averaged across mice. D) Mean motion activity during the 19 second pre-shock epoch for each of the trials. *Asterisks indicate significant differences between 10 and 60wo mice; two-way ANOVA with Tukey’s post hoc test.

**Supplemental Figure 7.** Averaged raw motion index traces over the course of day 2 of ‘fear recall’ illustrates consistency in response to conditioned stimulus (tone). Traces in arbitrary units were exported and normalized to allow qualitative comparisons. Blue arrows indicate the onset of tone. No unconditioned stimulus presented on day 2. A) Motion traces for 10wo mice. B) Motion traces for 30wo mice. C) Motion traces for 60wo mice. D) Mean motion activity during the 19 second tone epoch for each of the trials.

**Supplemental Figure 8.** Motion index activity averaged during the 4 second post tone onset period. A) Activity on day 1 of conditioning, B) Activity on day 2 of re-testing (‘recall’). Data are mean ± standard error. *Asterisks indicate significant differences, two-way ANOVA with Tukey’s post hoc test. **Asterisks indicate significant differences between 30 and 60wo mice; two-way ANOVA with Tukey’s post hoc test.

**Supplemental Figure 9.** Identified functional connectivity networks from pICA (see methods for details). Networks are qualitatively classified based on anatomical location of peak t statistic voxels (threshold free cluster enhancement corrected). Classification names and component order number are indicated overlying the statistical maps. Unilateral maps that are matched by a separate, but mirror image contralateral component is merged into a single map and both component numbers shown above maps. The right and left hemisphere sides are indicated at the bottom of the figure. Statistical maps overlaid onto multi-subject T2 anatomical template (t>2.3 shows significant voxels). N = 10/group

**Supplemental Figure 10.** Exploratory multivariate analyses between functional network measures and several affective behavior variables. A) Individual factor map with labeled individuals contributing to a high degree to the first two dimensions. B) Variables factor map with labeled variables projected best on the first two dimensions. Data points along the positive dimension 1 axis have high values for modularity and marginal distance activity. Data points in dimension 2 with high CPL values have low values for fear conditioning recall. Abbreviations in panel B: Assort: assortativity, CPL: characteristic path length, FCd1: fear conditioning day 1, FCd2: fear conditioning day 2, FCd3: fear conditioning day 3, Mod: modularity, MrgDist: marginal distance, SW: small world, Trans: transitivity.

**Supplemental Figure 11.** 2D scores plots of (**A**) principal component analysis (PCA) and (**B**) partial least squares discriminant analysis (PLS-DA) from GC-MS data of hippocampus region of the mouse brain. The variation is displayed in parentheses on each axis of PCA and PLS-DA. The shaded areas indicate 95% confidence regions. PLS-DA maximizes the separation among groups using the group label and the optimal number of components required to build the PLS-DA model was determined by cross validation. The robustness of the mathematical model was determined by sum of squares captured (R^2^) and the cross-validated R^2^ (Q^2^) parameters. Group sizes are: 10wo n = 4, 30wo n = 5, 60wo n = 6.

**Supplemental Figure 12.** 2D scores plots of (**A**) principal component analysis (PCA) and (**B**) partial least squares discriminant analysis (PLS-DA) from GC-MS data of prefrontal cortex (PFC) region of the mouse brain. The variation is displayed in parentheses on each axis of PCA and PLS-DA. The shaded areas indicate 95% confidence regions. PLS-DA maximizes the separation among groups using the group label and the optimal number of components required to build the PLS-DA model was determined by cross validation. The robustness of the mathematical model was determined by sum of squares captured (R^2^) and the cross-validated R^2^ (Q^2^) parameters. Group sizes are: 10wo n = 4, 30wo n = 5, 60wo n = 6.
