## Supplementary figures and images for "Age-Related Differences in Affective Behaviors in Mice: Possible Role of Prefrontal Cortical-Hippocampal Functional Connectivity and Metabolomic Profiles"

### Supplemental Figure 1

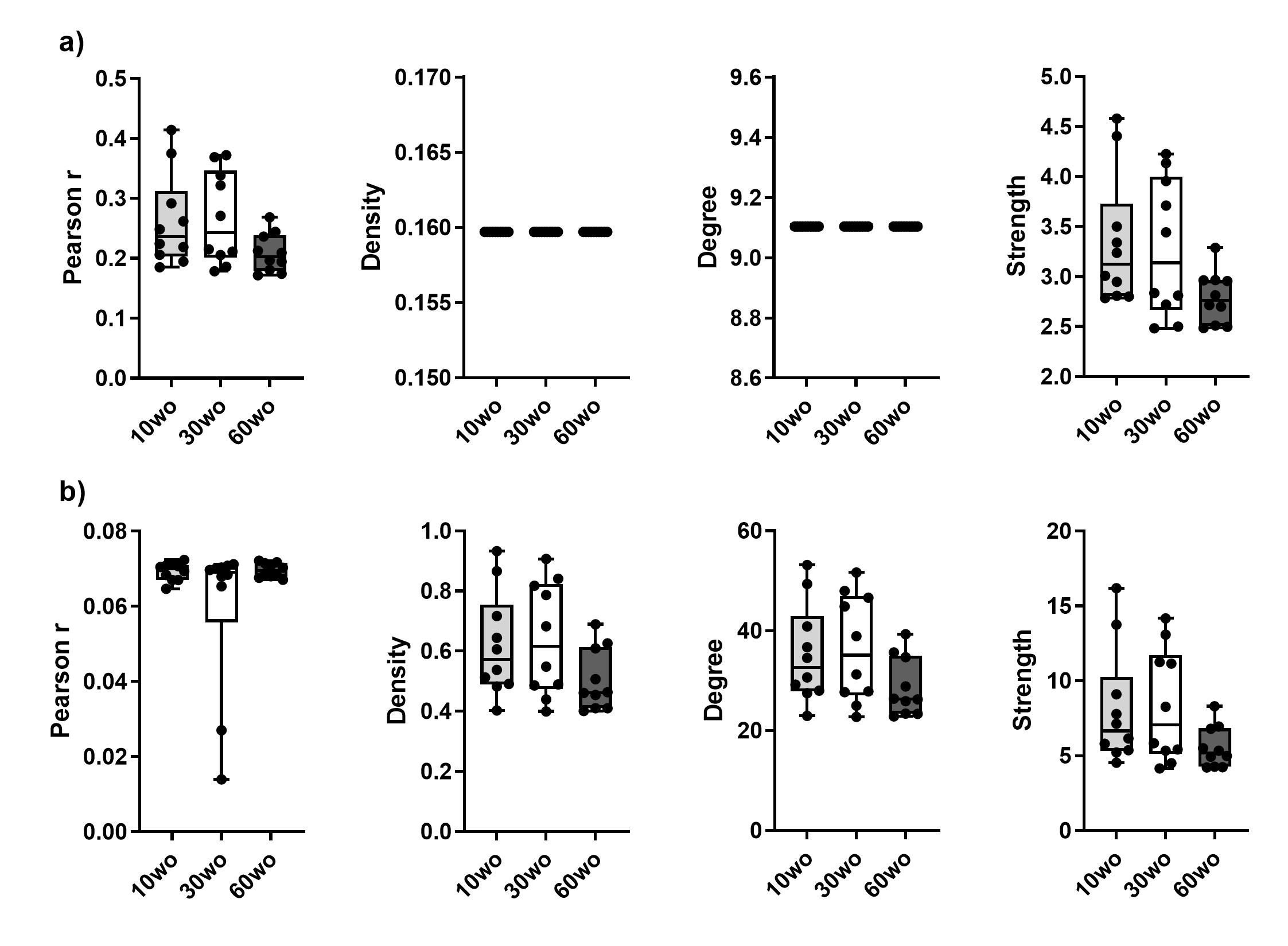

### Supplemental Figure 2

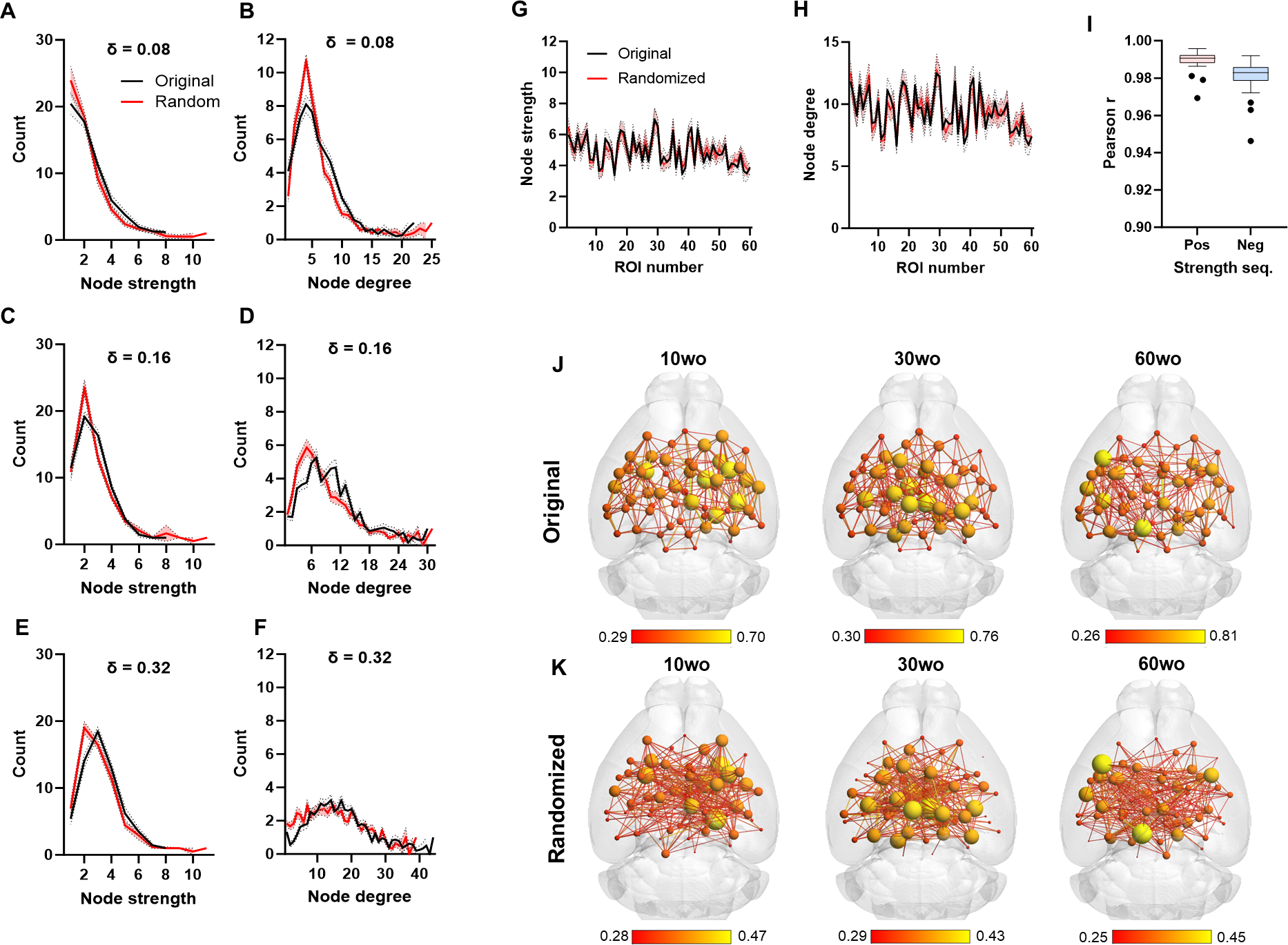

### Supplemental Figure 3

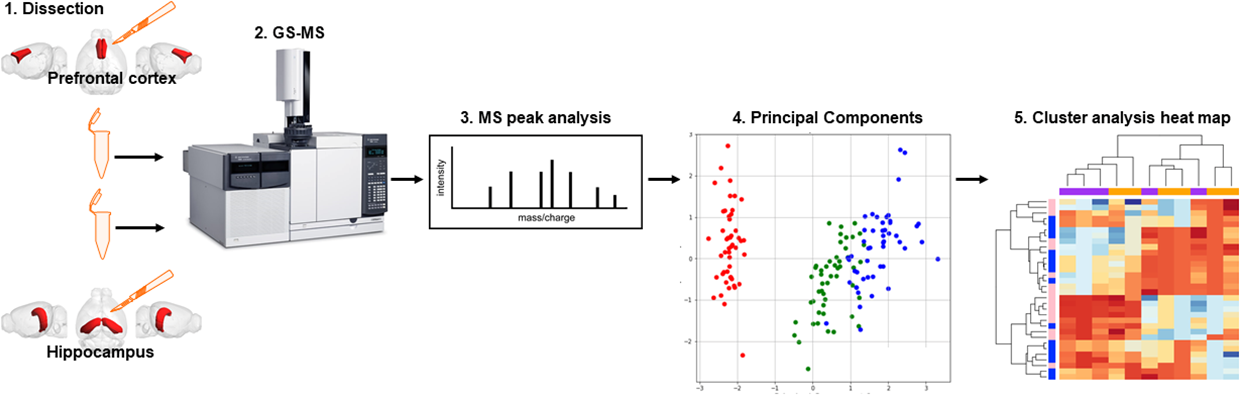

### Supplemental Figure 4

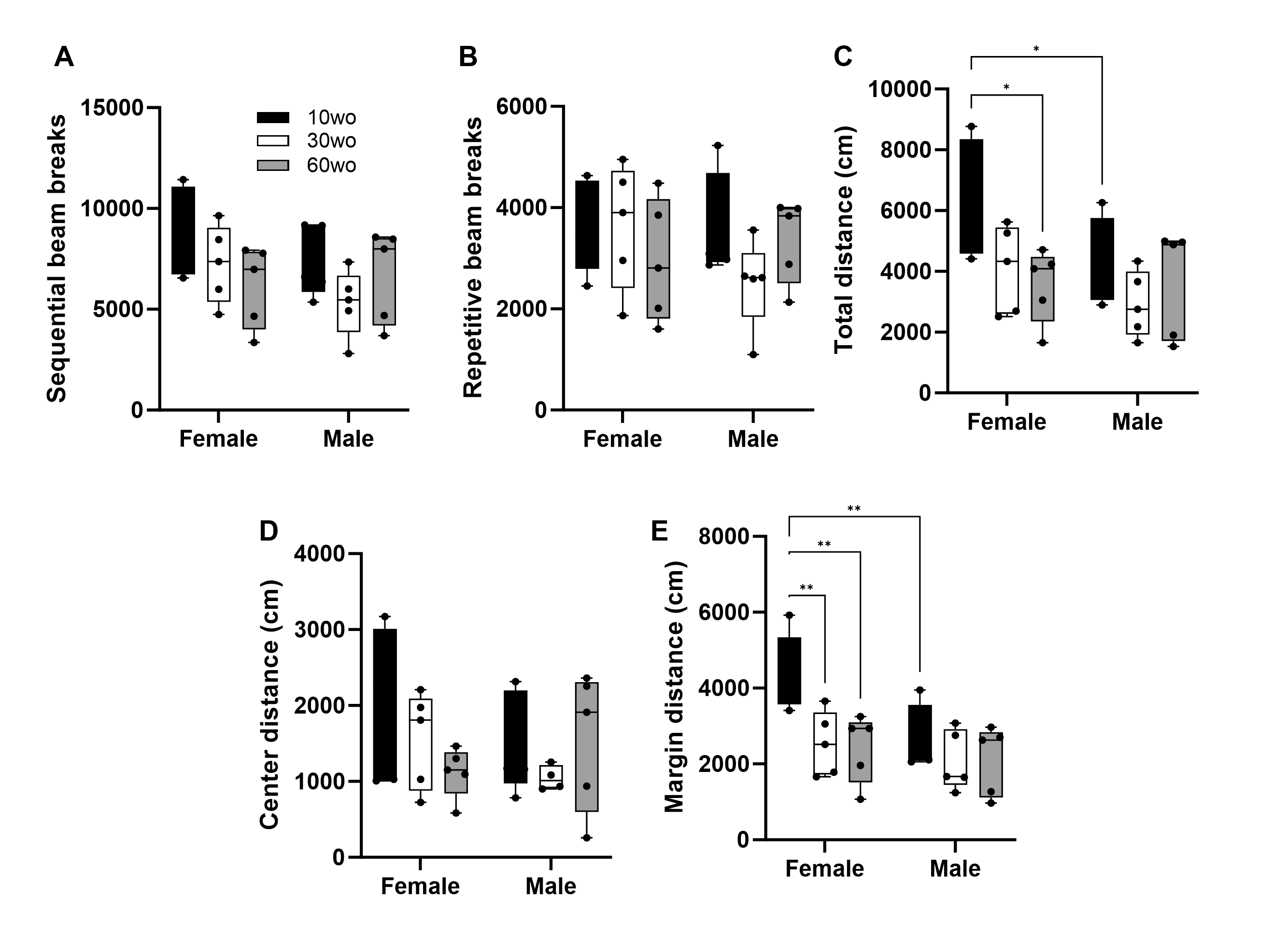

### Supplemental Figure 5

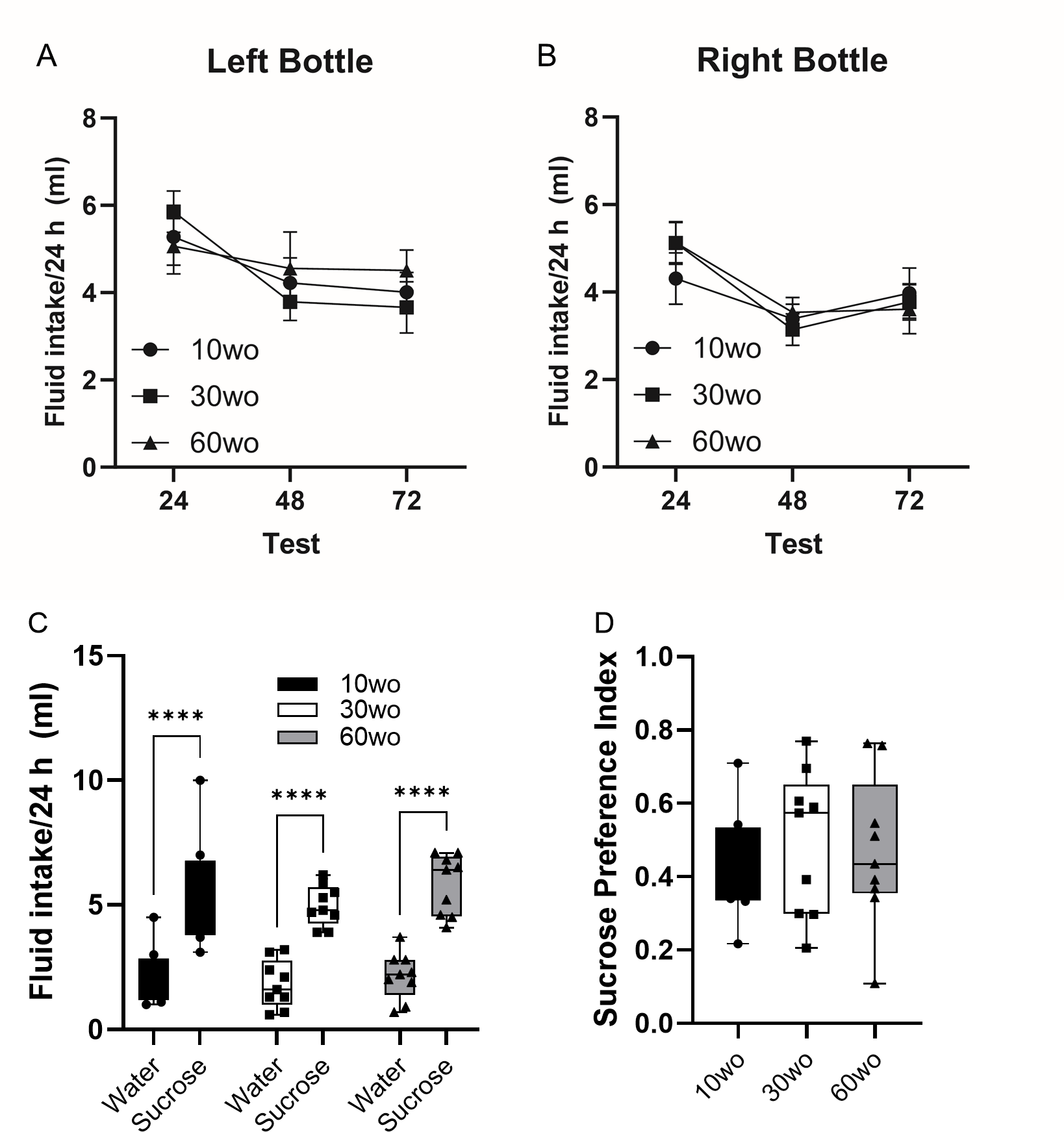

### Supplemental Figure 6

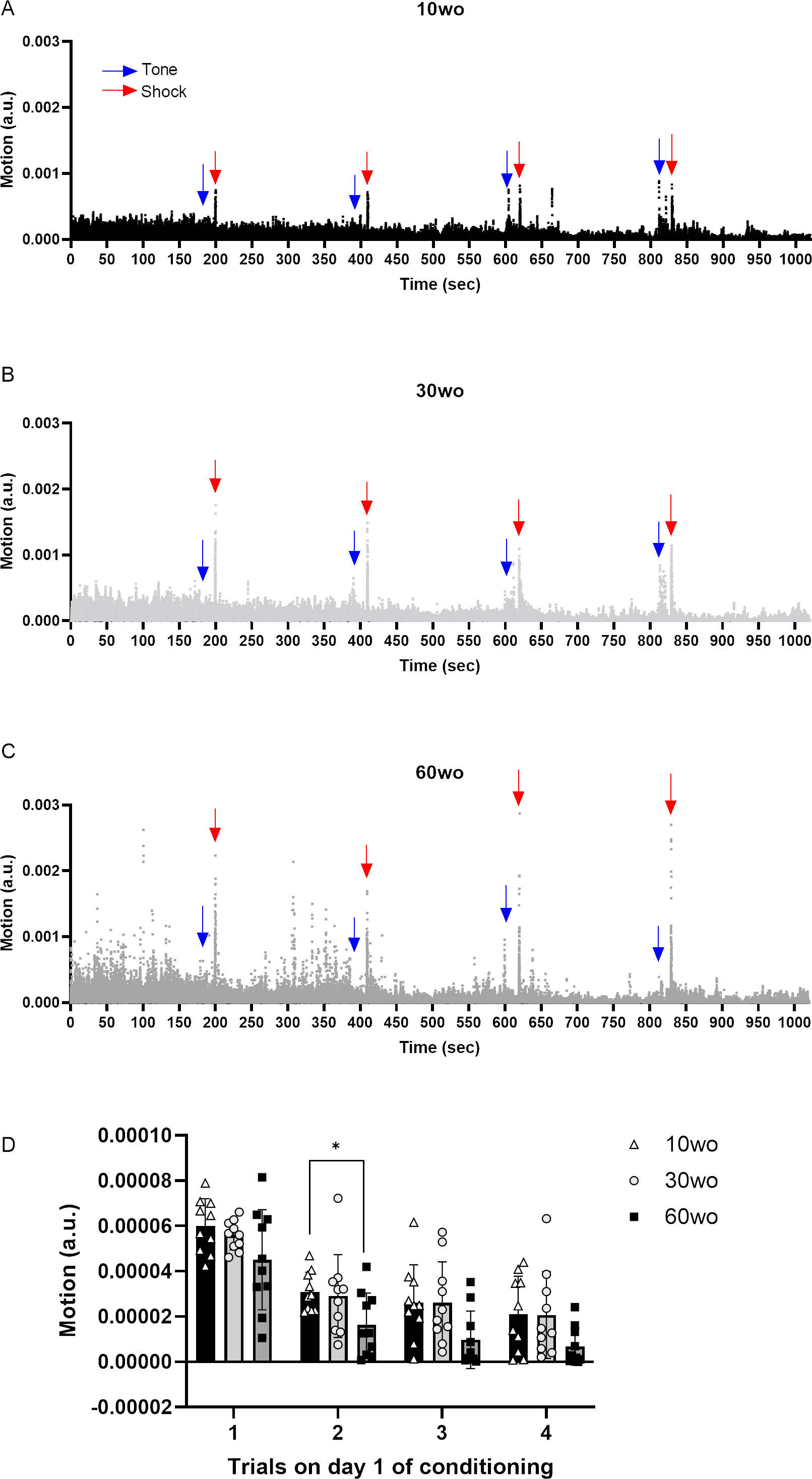

### Supplemental Figure 7

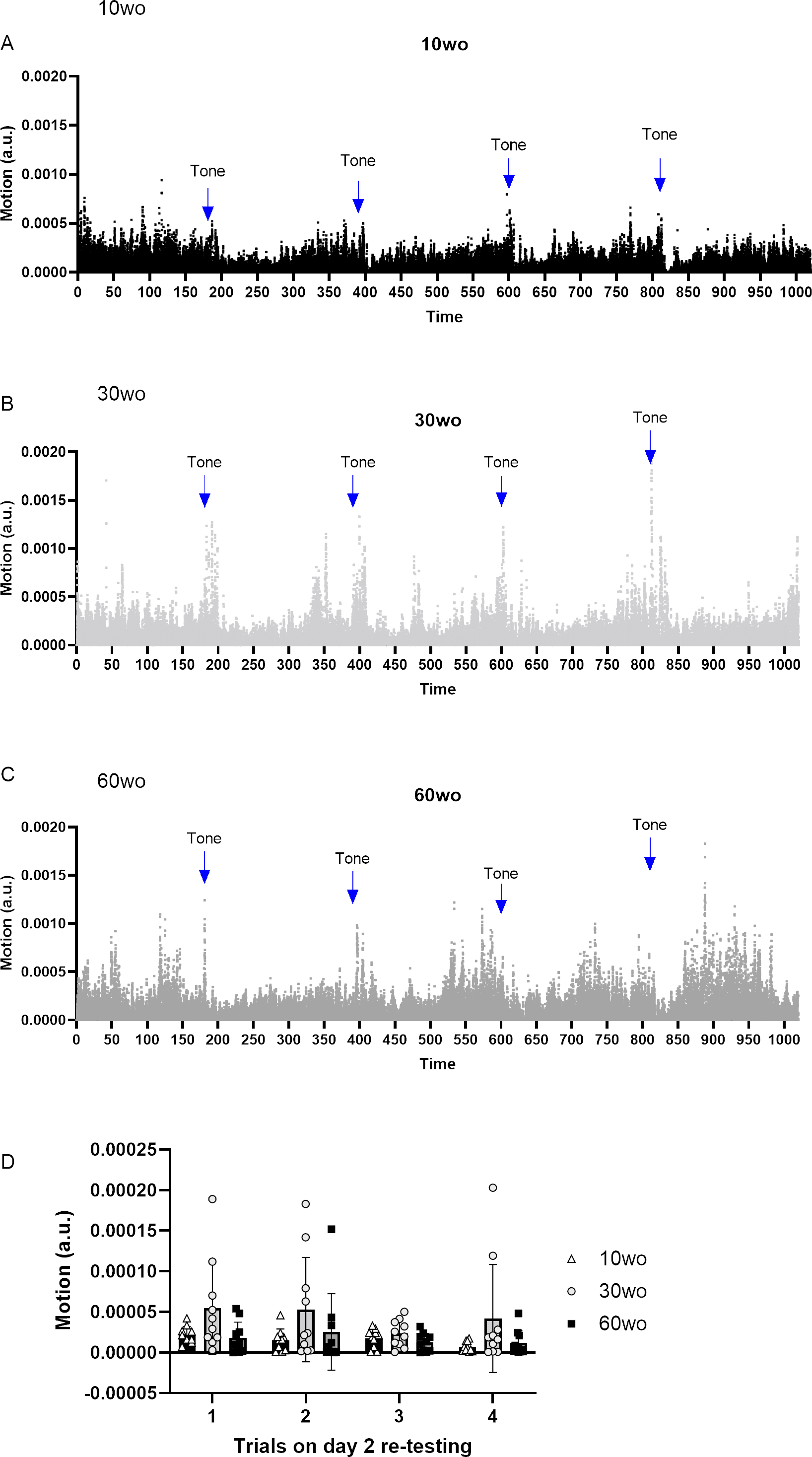

### Supplemental Figure 8

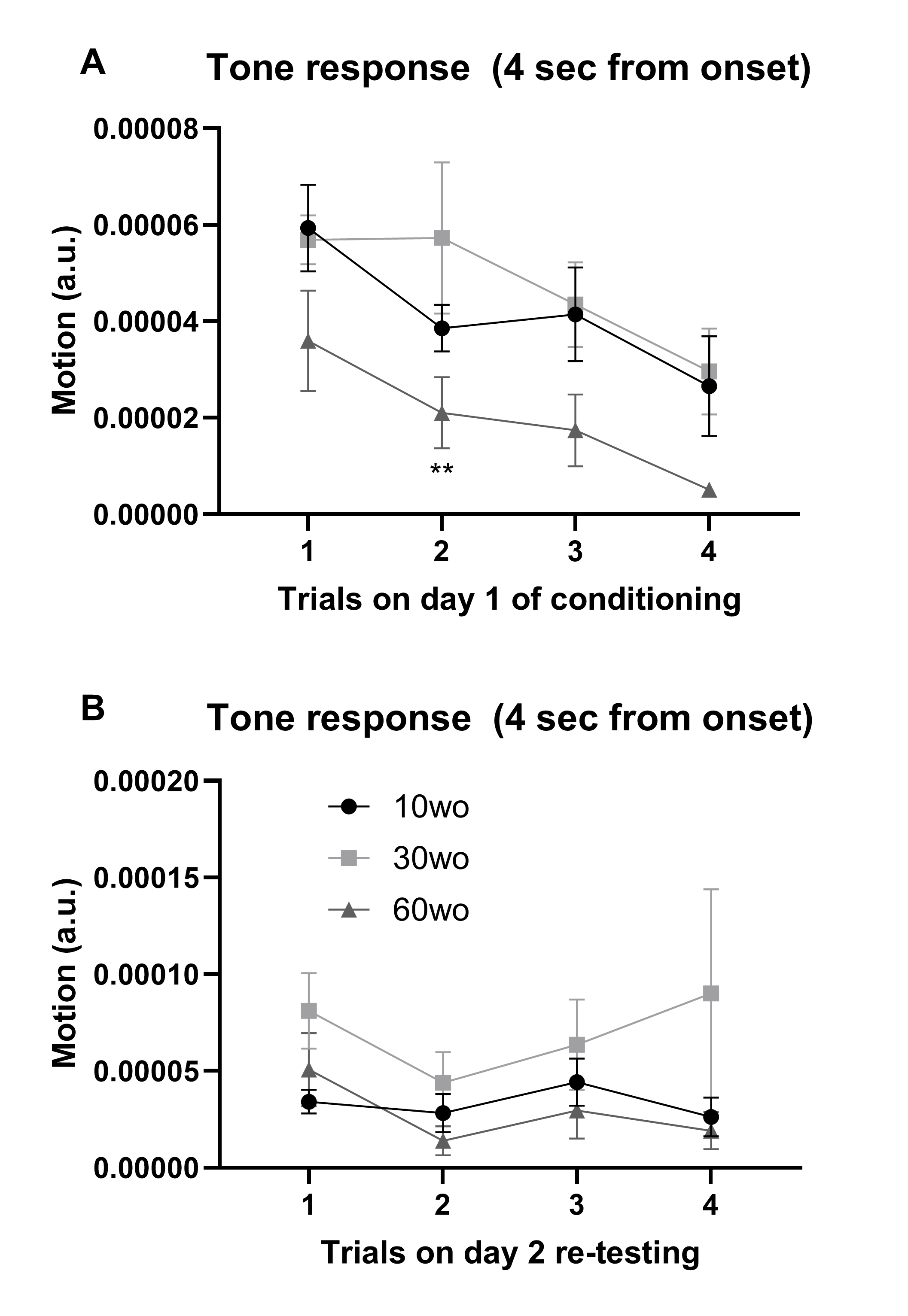

### Supplemental Figure 9

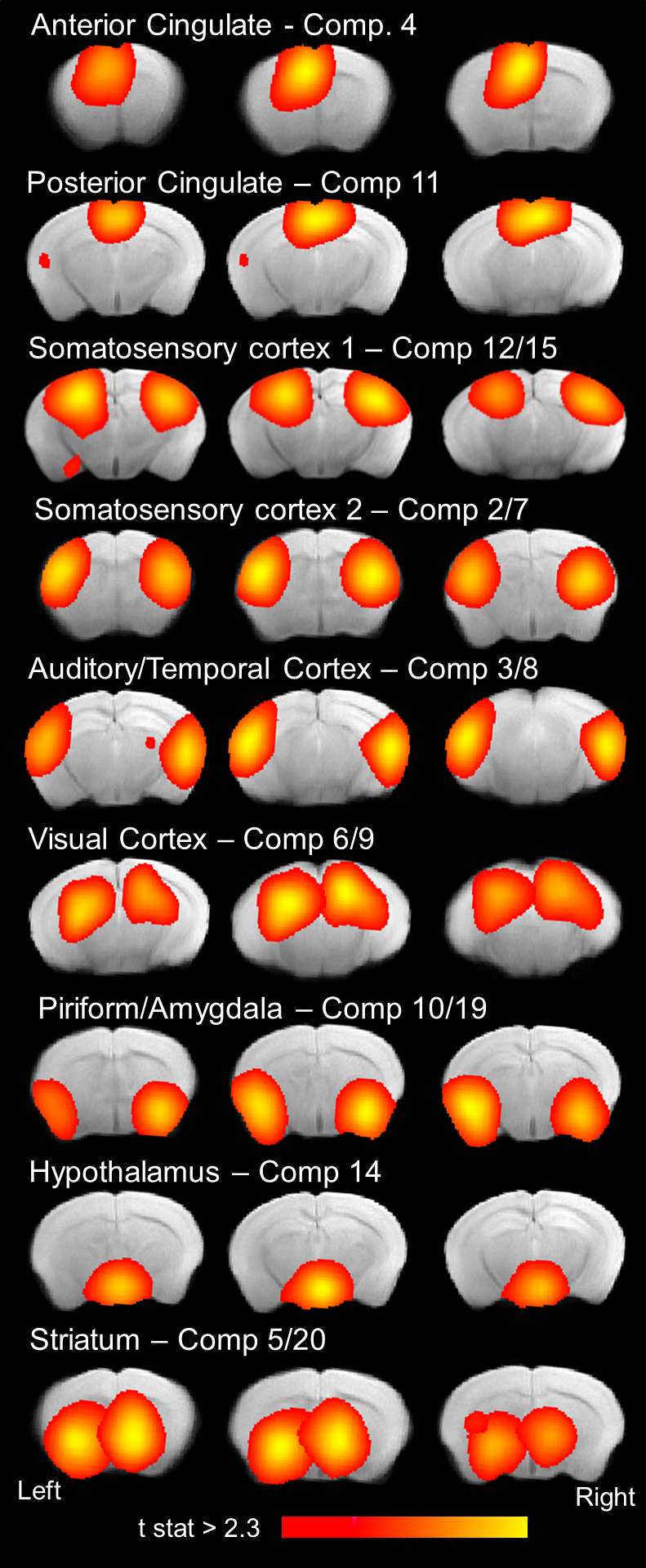

### Supplemental Figure 10

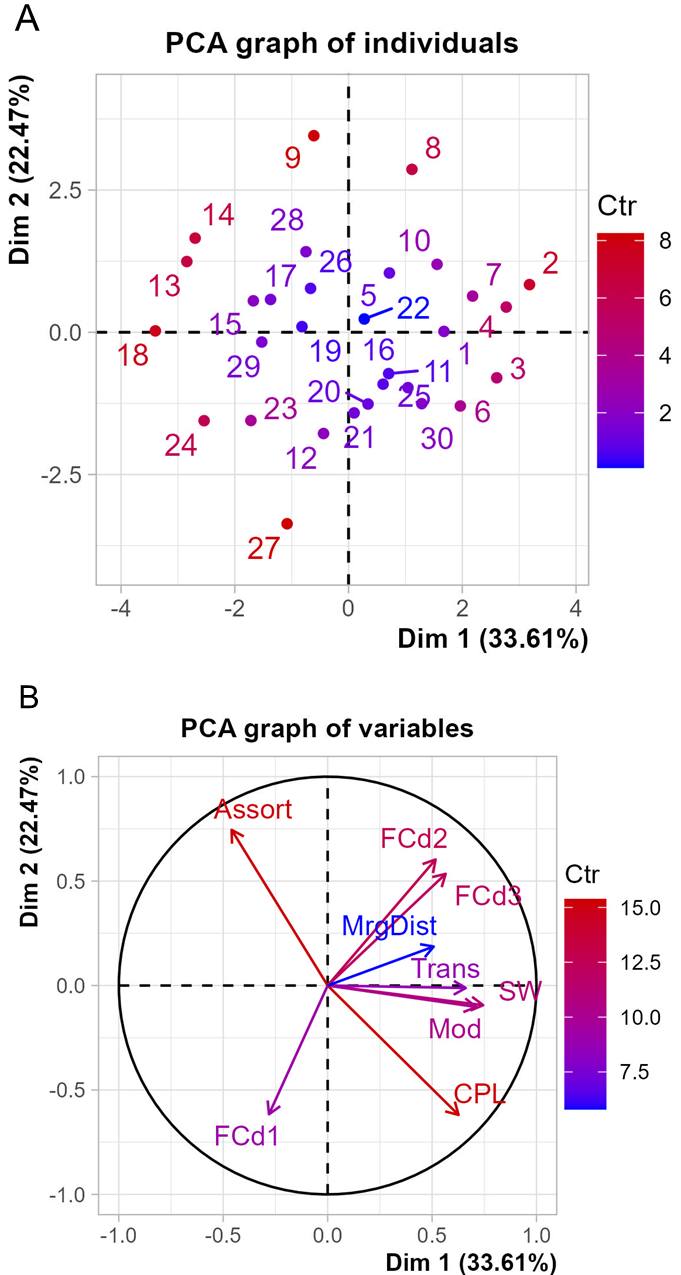

### Supplemental Figure 11

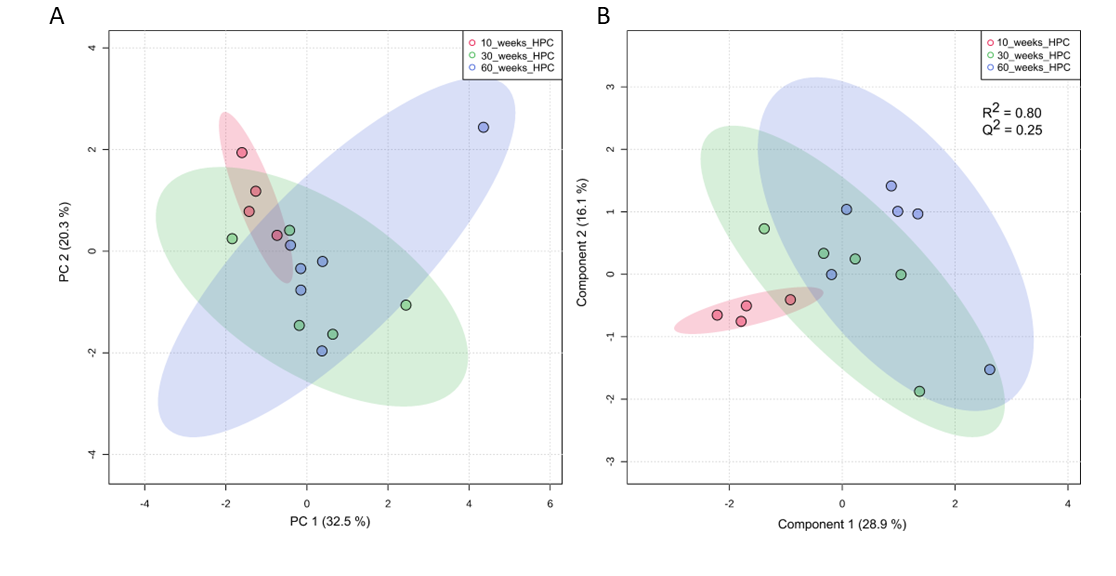

### Supplemental Figure 12

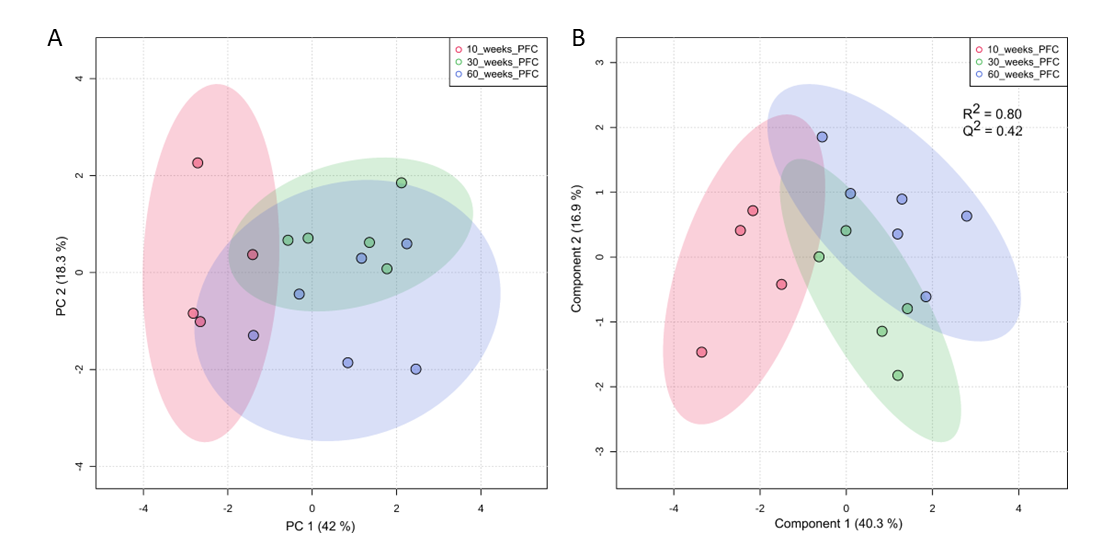
